# Computational pipeline provides mechanistic understanding of Omicron variant of concern neutralizing engineered ACE2 receptor traps

**DOI:** 10.1101/2022.08.09.503400

**Authors:** Soumya G. Remesh, Gregory E. Merz, Axel F. Brilot, Un Seng Chio, Alexandrea N. Rizo, Thomas H. Pospiech, Irene Lui, Mathew T. Laurie, Jeff Glasgow, Chau Q. Le, Yun Zhang, Devan Diwanji, Evelyn Hernandez, Jocelyne Lopez, Komal Ishwar Pawar, Sergei Pourmal, Amber M. Smith, Fengbo Zhou, QCRG Structural Biology Consortium, Joseph DeRisi, Tanja Kortemme, Oren S. Rosenberg, Anum Glasgow, Kevin K. Leung, James A. Wells, Kliment A. Verba

## Abstract

The SARS-CoV-2 Omicron variant, with 15 mutations in Spike receptor binding domain (Spike-RBD), renders virtually all clinical monoclonal antibodies against WT SARS-CoV-2 ineffective. We recently engineered the SARS-CoV-2 host entry receptor, ACE2, to tightly bind WT-Spike-RBD and prevent viral entry into host cells (“receptor traps”). Here we determine cryo-EM structures of our receptor traps in complex with full length Spike. We develop a multi-model pipeline combining Rosetta protein modeling software and cryo-EM to allow interface energy calculations even at limited resolution and identify interface side chains that allow for high affinity interactions between our ACE2 receptor traps and Spike-RBD. Our structural analysis provides a mechanistic rationale for the high affinity (0.53 - 4.2nM) binding of our ACE2 receptor traps to Omicron-RBD confirmed with biolayer interferometry measurements. Finally, we show that ACE2 receptor traps potently neutralize Omicron- and Delta-pseudotyped viruses, providing alternative therapeutic routes to combat this evolving virus.

## Introduction

The rapidly evolving SARS-CoV-2 virus has accumulated several mutations throughout the pandemic. The Omicron (B.1.1.529) variant was first reported in November 2021 in South Africa to have 37 mutations in its Spike glycoprotein and was quickly designated as a variant of concern (VOC) by the World Health Organization (Viana et al., 2022; Walls et al., 2020). The Omicron Spike (S) glycoprotein receptor-binding domain (RBD) and N-terminal domain (NTD) harbor 15 and 11 mutations, respectively, leading to lower plasma neutralization in patients previously infected with other SARS-CoV-2 variants or in fully vaccinated individuals (Cameroni et al., 2022; Cao et al., 2022; Cele et al., 2021; Liu et al., 2022; Mannar et al., 2022; Planas et al., 2022; VanBlargan et al., 2021; Wilhelm et al., 2021). Due to the antigenic shift in the Omicron variant, currently only two out of eight clinical monoclonal antibody treatments, S309 (sotrovimab parent) and the COV2-2196/COV2-2130 cocktail (cilgavimab/tixagevimab parents), retain appreciable neutralizing capacity albeit reduced by 2-3 and 12-200 fold, respectively compared to neutralization of Wuhan-hu-1 strain (Cameroni et al., 2022; Cao et al., 2022; Liu et al., 2022; Mannar et al., 2022; Planas et al., 2022; VanBlargan et al., 2021). An earlier VOC, the B.1.617.2 Delta variant, also acquired 10 mutations in the Spike S glycoprotein, outcompeted other circulating virus isolates, and enhanced transmission and pathogenicity while diminishing antibody-based neutralization activity (McCallum et al., 2021; Mlcochova et al., 2021). Interestingly, both the Delta- and Omicron RBD continue to bind the SARS-CoV-2 entry receptor, human angiotensin converting enzyme 2 (ACE2), with almost 2-fold higher affinity than the wild-type Spike-RBD (Cameroni et al., 2022; Mannar et al., 2022). Previously, we developed engineered ACE2 “receptor traps” as viable candidates for SARS-CoV-2 virus neutralization (Glasgow et al., 2020). The receptor traps were computationally designed and further affinity-optimized by yeast display. The optimized ACE2 extracellular domains were fused to a human IgG1 Fc domain to afford additional binding avidity and neonatal Fc receptor (FcRN) binding for long circulating half-life. One advantage of using an ACE2 receptor trap as a therapeutic to treat SARS-CoV-2 infections is that resistance evolved to ACE2 traps would also likely render the virus unable to infect host cells via the ACE2 entry receptors. Thus, ACE2 traps could provide alternative therapeutic strategies for the rapidly evolving SARS-CoV-2 virus.

Our computationally designed (CVD293) and affinity matured (CVD313) ACE2 Fc-fusions have ∼20-25-fold improved virus neutralization ability against WT-SARS-CoV-2 pseudotyped lentivirus (Wuhan-hu-1 strain) compared to the WT-ACE2 Fc-fusion (Glasgow et al., 2020). However, the structures of neither CVD293 nor CVD313 bound to WT-Spike-RBD have been captured, leaving it an open question as to why the affinity matured ACE2 receptor traps bind WT-Spike-RBD tighter. Here, we shortened the length of the linker between the ACE2 extracellular domain (residues 18-740) of CVD313 and the Fc domain to ∼13 amino acids to generate the construct CVD432, which has an improved mammalian expression profile (total yield post purification of CVD432 is ∼2.5 times more than CVD313). We determined the cryo-EM structures of wild-type full-length Spike protein (WT-fl-Spike) with either CVD293 or CVD432, which revealed molecular details of the interactions between the ACE2 receptor traps and fl-Spike. Building on previous approaches to provide ensemble models from cryo-EM maps (Herzik et al., 2019), we developed a multi-model workflow to improve our confidence in the molecular interactions and the interaction energies of the ACE2/RBD. This analysis allowed us to detect subtle interface differences between the two receptor traps that were critical for the improved ACE2/RBD interface. Furthermore, we used the cryo-EM structures together with Rosetta interface energy calculations to model the interactions between our ACE2 receptor traps and Spike-RBD of Omicron VOC of SARS-CoV-2, rapidly identifying the direct contact residues and predicting an even tighter interaction than to the Spike-RBD of Wuhan-hu-1 strain. We validated these structural modeling results by performing biolayer interferometry measurements. Finally, we showed that our ACE2 receptor traps potently neutralize Delta- and Omicron-SARS-CoV-2 pseudotyped viruses and thus can serve as alternate therapeutic candidates against SARS-CoV-2 infections.

## Results

### Cryo-EM reconstructions of WT-fl-Spike trimer in complex with engineered ACE2 receptor traps

To understand the molecular details of the interactions between the WT-fl-Spike and the engineered ACE2 receptor traps CVD293 and CVD432, we determined the cryo-EM structures of these complexes (Figure 1). We confirmed that both CVD293 and CVD432 potently neutralize WT-SARS-CoV-2 (Wuhan-hu-1 B.1 strain with D614G mutation only) pseudotyped virus (Figure S1d). As noted previously for the WT-fl-Spike/WT-ACE2 complex (Yan et al., 2021; Zhou et al., 2020a), we observed conformational heterogeneity for the complexes between WT-fl-Spike and the engineered ACE2 receptor traps, with the number of RBDs in the “up” or “down” conformation per spike protein varying in the ACE2-bound state. The complex between WT-fl-Spike and CVD293 showed a 1-RBD-up WT-fl-Spike with full ACE2 occupancy (∼20%), an appreciable percentage of 1-RBD-up WT-fl-Spike with partial ACE2 occupancy (∼54%), a 2-RBD-up state with 1-ACE2 occupancy (∼15%) and a 1-RBD-up state with no ACE2 occupancy (Figure 1a) per trimer. On the other hand, WT-fl-Spike and CVD432 showed a 1-RBD-up WT-fl-Spike with full ACE2 occupancy (∼12%), a 2-RBD-up state with 2-ACE2 occupancy (∼9%), all-RBD-down state, and other partial- to no-ACE2 occupancy 1-RBD- and 2-RBD-up states (Figure 1b). While ACE2 residues 18-614 were well resolved and could be fit into our cryo-EM maps of the complexes between WT-fl-Spike/ACE2 receptor traps, we could not model the highly flexible collectrin domain of the ACE2 traps (residues 615-740) as well as the Fc-domains. Nevertheless, the WT-Spike-RBD/ACE2 receptor trap sub-region could be resolved at low to medium resolution.

**Figure 1.**
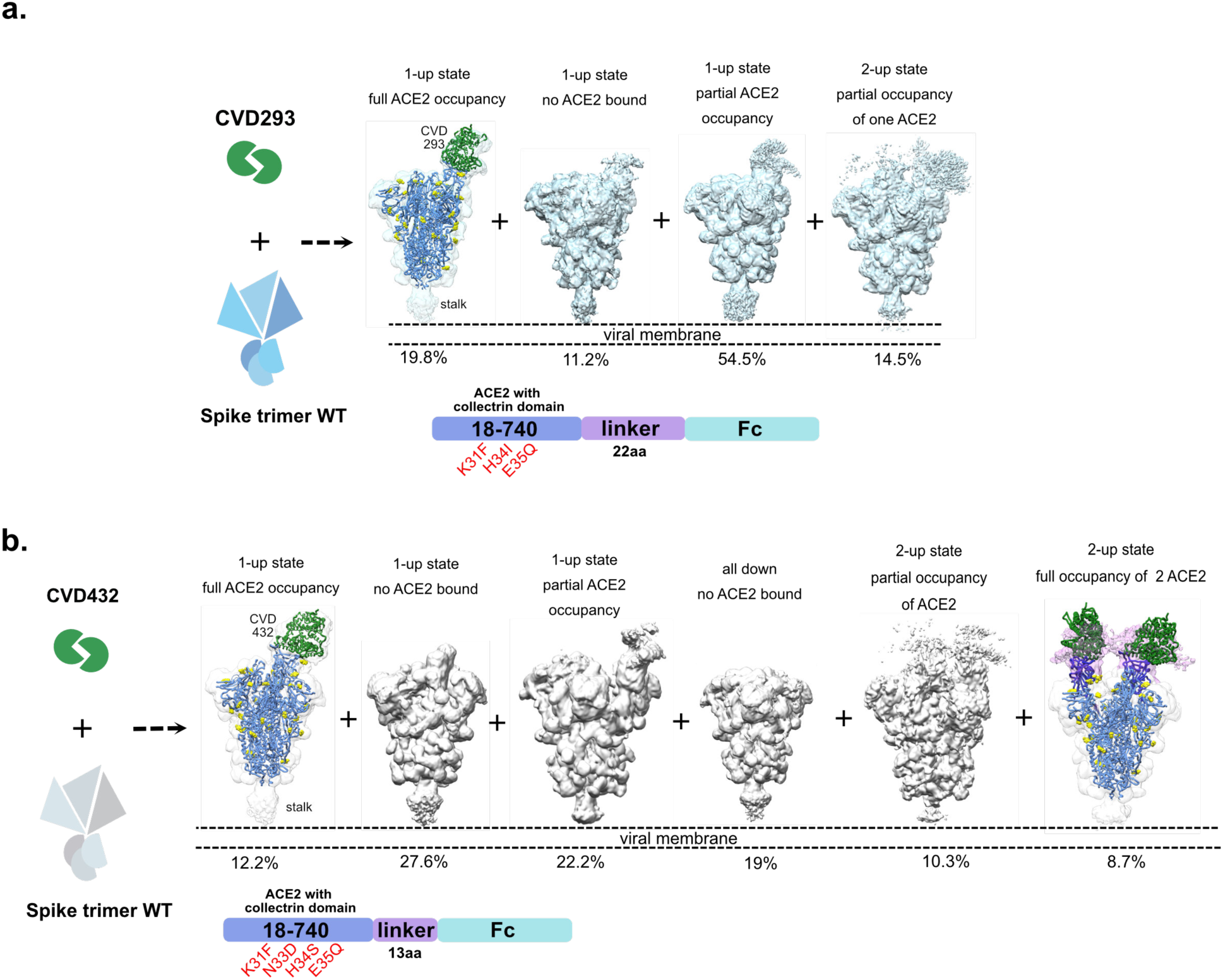
Cryo-EM reconstruction of WT-fl-Spike with computationally designed, CVD293 or linker variant of the affinity matured variant, CVD432. **a-b**. Cryo-EM reconstructions of WT-fl-Spike with CVD293 or CVD432 showing the heterogeneity in distribution of all RBD down, 1-RBD- or 2-RBD-up states and variable ACE2 occupancy. Also shown is schematic of the primary structure of CVD293 or CVD432 and the engineered mutations, colored by domain.

Local refinement of CVD293 or CVD432 (residues 18-614) with WT-Spike-RBD (residues 330-541, Wuhan-hu-1 strain) generated maps of resolution ∼3.8-4.8 Å and 3.4-3.8 Å, respectively, for the interface residues (Figure 2a, S2, S3, S4 and S5). As described above, the high level of heterogeneity between the 1-RBD-up and 2-RBD-up states for WT-fl-Spike with either CVD293 or CVD432 made obtaining high resolution maps of the interface residues challenging. To better understand the modalities of continuous protein motions within the 1-RBD-up or 2-RBD-up states, we utilized 3D variability analyses (3DVA) (Punjani and Fleet, 2021) (Figure S1a-c). Both WT-fl-Spike/CVD293 and WT-fl-Spike/CVD432 complexes showed considerable rotation relative to the vertical axis (∼5-7°) in the 1-RBD-up/ACE2 bound sub-region of the cryo-EM map (Figure S1a, b). In contrast, the 2-RBD-up state of WT-fl-Spike/CVD432 showed lateral shift of the ACE2 bound RBD sub-region of the cryo-EM map relative to the vertical axis of rotation (Figure S1c). This high degree of variability in the ACE2 bound/RBD sub-region of the cryo-EM map further impeded high resolution structure determination of the interface residues. Consequently, the precise rotamer positions, especially of key interface residues for CVD293 (engineered mutations - K31F, H34I, E35Q) or CVD432 (engineered mutations - K31F, N33D, H34S, E35Q) could not be obtained with high confidence directly from the cryo-EM maps. (Figure 2b, c; cryo-EM consensus models are shown fit to the cryo-EM maps).

**Figure 2.**
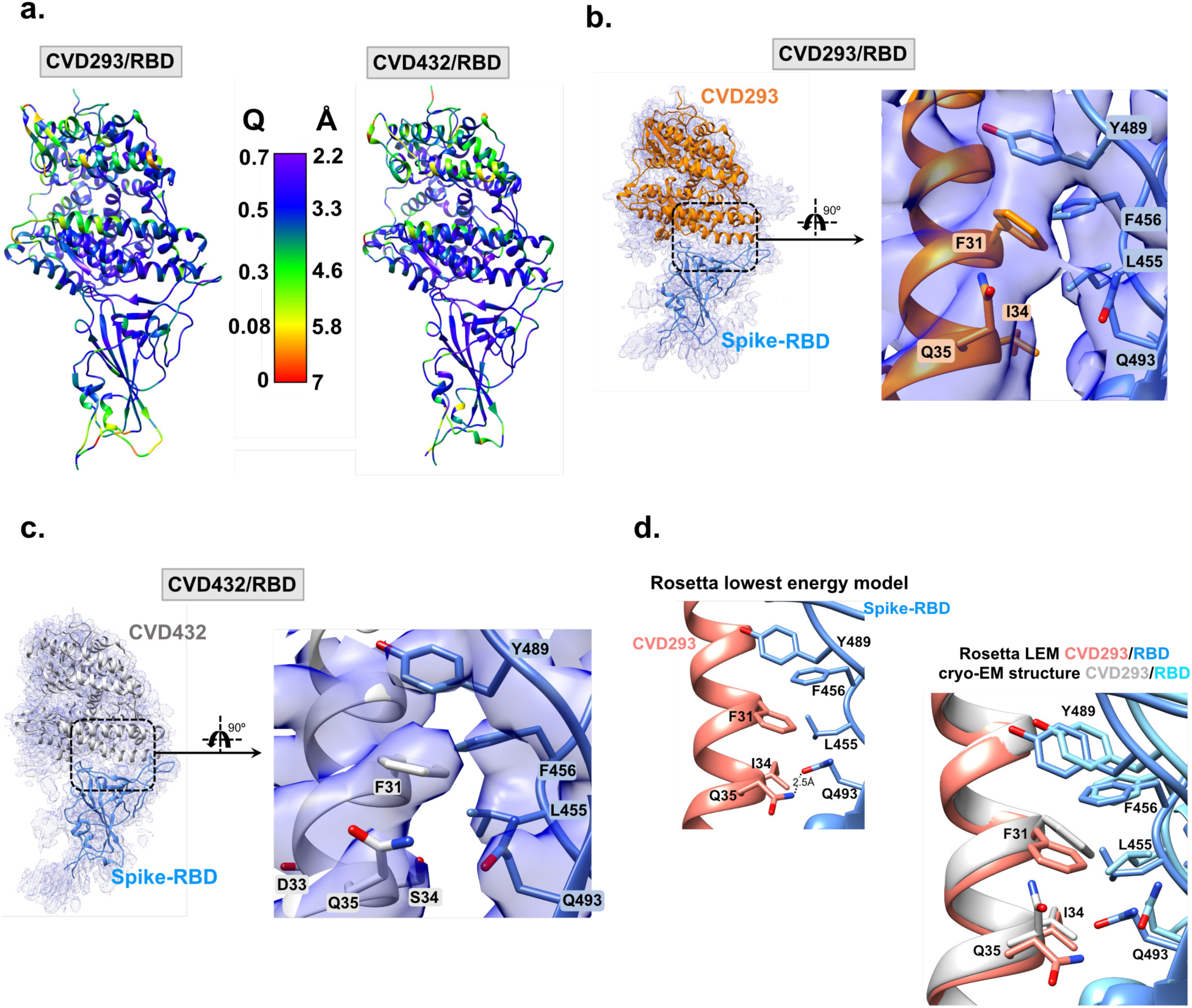
Cryo-EM reconstruction of WT-Spike-RBD with engineered ACE2 Fc-fusions reveal contributions from hydrophobic interactions at RBD-ACE2 interface. **a**. WT-Spike-RBD/CVD293 and WT-Spike-RBD/CVD432 models colored by estimated per residue Q-score ranging from 0 (red) to 0.7 (purple). The color bar shows corresponding estimated resolution in Å for each Q-score. Expected Q-score for 3.5 Å map is 0.49 and expected Q-score for 3.36 Å map is 0.52. **b-c**. Cryo-EM reconstructions of WT-Spike-RBD with either CVD293 or CVD432 show favorable π–π stacking interactions between WT-Spike-RBD residue Y489 and engineered ACE2 residue F31. Additionally, there are also hydrophobic interactions between WT-Spike-RBD residue L455 and CVD293 residue I34 are also seen. Hydrogen bond interactions between WT-Spike-RBD residue Q493 and CVD293 or CVD432 residue Q35 are not apparent in the cryo-EM consensus model. **d**. The Rosetta lowest energy model for CVD293 is overlaid with the cryo-EM model. Both models show hydrophobic and hydrogen bond interactions between CVD293 and WT-Spike-RBD residues that contribute to improved interface energy (REU) compared to the ACE2-WT Spike RBD interaction.

### Improving confidence of the interface interactions in limited resolution cryo-EM maps through Rosetta enabled multi-model workflow

Inspired by a previous ensemble model refinement approach to improve confidence in maps with resolution variation (Herzik et al., 2019), we developed a multi-model workflow which allowed for calculation of an “average predicted interface energy” metric across an ensemble of models consistent with the cryo-EM map in addition to the previously reported “average RMSD” metric. Together, these metrics provided a statistics-based view of the ACE2/RBD interface (Figure 3a). Briefly, 10-residue overlapping stretches of the ACE2 interface helix (residues 21 - 52) of each cryo-EM consensus model (WT-Spike-RBD (330-541)/CVD293 (18-614) or WT-Spike-RBD (330-541)/CVD432 (18-614)) were subjected to a CartesianSampler mover within Rosetta that samples similar sequence and secondary structure fragments from within PDB and locally minimizes them into the cryo-EM density, generating 2000 models for each 10-residue stretch. Each model was then all-atom minimized within the cryo-EM map using FastRelax mover with a scoring term for the model agreement with the cryo-EM map, as previously described (DiMaio et al., 2015). The Rosetta parameters and scoring functions used were based on the estimated map resolution (Wang et al., 2016). This refinement protocol was iteratively run to generate a total of ∼8000 overlapping atomic models of the interface helix.

**Figure 3.**
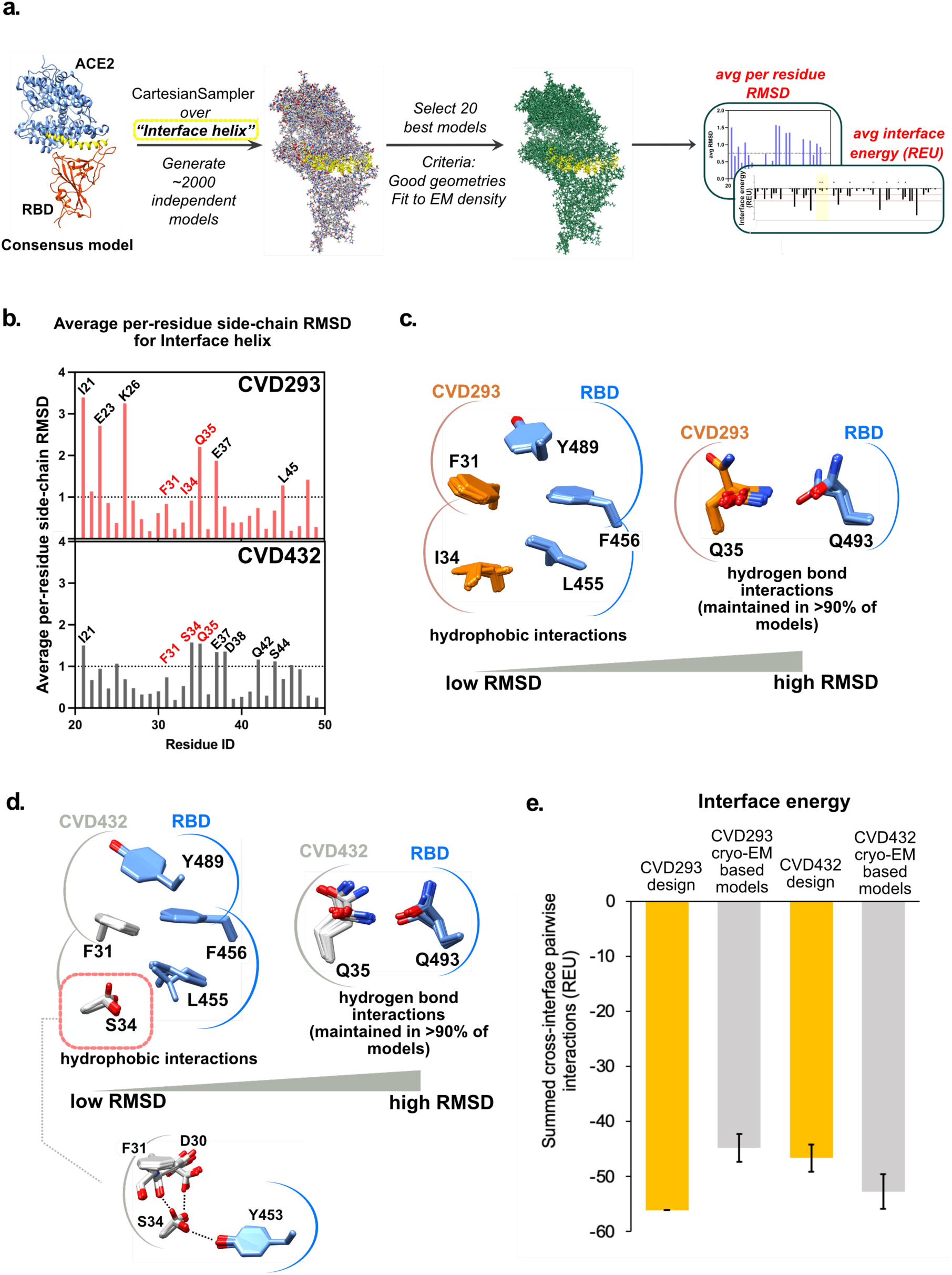
Multi-model pipeline improves confidence of molecular interactions at the interface residues in cryo-EM derived models of WT-Spike-RBD with engineered ACE2 Fc-fusions. **a.** Multi-model pipeline with average Rosetta interface energy and average per-residue side-chain RMSD metrics for interface residue rotamer positions. **b**. Average per-residue side chain RMSD for interface helix residues of CVD293 and CVD432 **c-d.** Superposition of critical interface residues of the top 80 selected cryo-EM based models for CVD293 and CVD432. **e**. Average Rosetta interface energy for CVD293 design model, CVD293 cryo-EM based models, CVD432 design model and CVD432 cryo-EM based models.

We next ranked the atomic models based on total Rosetta scores ensuring good geometries (pick top 200) and fits to the cryo-EM map (pick top 20 out of the above 200) for each 10-residue stretch to select a total of 80 models for the entire interface helix. The average per residue side-chain RMSDs and predicted average interface energy for all the residues of the interface helix for the top selected cryo-EM based models were then calculated (Figure 3b,e). We compared average per-residue side-chain RMSDs as we expected very small changes in the average backbone RMSDs which can cause side-chain discrepancies being down-weighted in average full-residue RMSDs. We superimposed the interface helix residues (residues 21-52) of the top 80 selected cryo-EM based models for both CVD293 and CVD432 to analyze the convergence of the side-chain conformations and the intermolecular interaction with corresponding WT-Spike-RBD residues. We observed that while the interface residues of CVD293 (F31, I34) or CVD432 (F31) with low average side-chain RMSD make well-defined hydrophobic interactions with corresponding residues in WT-Spike-RBD (L455, F456, Y489), the high average side-chain RMSD residue (Q35 of both CVD293 and CVD432) makes hydrogen bond interactions with neighboring WT-Spike-RBD residue (Q493) in over 90% of the atomic models (Figure 3c, d). We further noted that CVD432 high average side-chain RMSD residue, S34, can make both inter- (with WT-Spike-RBD residue Y453) and intra-molecular hydrogen bonds (with backbone carbonyl atoms of residues D30 or F31). We infer that the low average side-chain RMSD engineered hydrophobic residues of the ACE2 receptor traps likely provide the key functional interactions responsible for the improved binding affinity of the engineered receptor traps for WT-Spike-RBD.

### Analysis of the cryo-EM based models of CVD293 (18-614) and CVD432 (18-614) in complex with the WT-Spike-RBD (330-541)

The comparison between the original computationally designed model of the CVD293/WT-Spike-RBD complex and the cryo-EM based models of the complex determined with our multi-model workflow revealed high overall structural agreement (Figure 2d). The average Cα-RMSD between the computationally designed model and the experimentally solved cryo-EM consensus model of CVD293 (residues 18-614) was 0.93 Å, and the average Cα-RMSD for the interface helix (ACE2 residues 21-88, 322-329, 352-358 and Spike residues 444-446, 475-505) was 0.41 Å.

We next compared the calculated interface energy for the CVD293/WT-Spike-RBD interface in the design model and the cryo-EM based models. We found that the predicted interface energy is considerably lower for the CVD293 design model (-58 REU) than the average of 80 CVD293 cryo-EM based models (-45 REU) (Figure 3e). This discrepancy between the predicted interface energy for the design model and the average interface energy calculated from the 80 cryo-EM based models is likely due to several differences in side-chain-mediated interactions involving designed residues. For example, isoleucine 34, a designed residue in CVD293, is predicted to contribute less to the interface energy in the cryo-EM based models (average from 80 atomic models of CVD293: -2.5 REU) than in the design model (-3 REU), despite strong agreement in the atomic coordinates for this residue across all cryo-EM based models (Figure 2d). The predicted interaction energy of I34 is worse in the cryo-EM models than the design model because its interaction partner across the interface, residue L455 of WT-Spike-RBD, universally adopts a different rotamer than the one in the original design model (Figure S7a).

Residue Q35, another designed residue in CVD293, showed the largest observed average per-residue side-chain RMSD among the designed positions on ACE2 in the cryo-EM based models (Figure 3b). Although the Q35 side-chain often occupies a different rotamer in the cryo-EM based models than in our original design model of CVD293, it still makes a hydrogen bond with the WT-Spike-RBD residue Q493 as intended (Figure 3c). The WT-Spike-RBD residue Q493 shows minimal conformational heterogeneity across the 80 cryo-EM based models of the complex CVD293/WT-Spike-RBD, and it makes intramolecular hydrogen bonds with the carbonyl group of the WT-Spike-RBD residue F490 to maintain the same loop conformation as seen in the complex between WT-ACE2 and WT-Spike-RBD (Figure S7a). It is plausible that the minimal average side-chain RMSD of residue Q493 and the overall similar loop conformation around residues F490-Q493 in the complex CVD293/WT-Spike-RBD contribute to the observed conserved hydrogen bond with residue Q35.

Another WT-ACE2 residue, K31, was mutated to a phenylalanine in CVD293. Consequently, the hydrogen bond between the WT-ACE2 residue K31 and the WT-Spike-RBD residue Q493 was lost in the complex CVD293/WT-Spike-RBD. Instead of making this hydrogen bond, the designed residue F31 in CVD293, with less than 1 Å average observed average side-chain RMSD, makes good packing interactions with other aromatics at the WT-Spike-RBD/engineered ACE2 trap interface as expected (Figure 3c). Overall, the average Cα-RMSD between the computationally designed model and the experimentally solved cryo-EM consensus model of CVD293 (residues 18-614) is close to 1 Å. However, using the cryo-EM based models of complex CVD293/WT-Spike-RBD determined with our multi-model workflow, specific residue-pair interactions that were critical to improving the WT-ACE2/WT-Spike-RBD interface with our computational design strategy are revealed.

We next wanted to understand why the affinity maturation of CVD293 in our yeast surface display campaign resulted in the I34S interface mutation in the construct CVD432, which has improved binding affinity for WT-Spike-RBD. We first generated a CVD432 “design” model by mutating the residue I34 in the CVD293 cryo-EM consensus model to a serine *in silico*. We also made an N33D mutation to CVD293 *in silico* even though this position is outside the RBD binding interface, because this mutation was also identified by yeast surface display in CVD432. The calculated interface energy for the CVD432 design model/WT-Spike-RBD interface is higher (-47 REU) than the average interface energy from 80 cryo-EM based models of the complex CVD432/WT-Spike-RBD (-53 REU) (Figure 3e). It can be inferred that simple *in silico* mutations do not provide a full explanation for the improved binding affinity between the engineered ACE2 trap and the WT-Spike-RBD, highlighting the value of the cryo-EM structure solution.

Comparing the cryo-EM based models of the complexes CVD293/WT-Spike-RBD and CVD432/WT-Spike-RBD, we find that the calculated interface energy from the average of 80 cryo-EM based models of the complex CVD432/WT-Spike-RBD is lower (-53 REU) than the average of 80 cryo-EM based models of the complex CVD293/WT-Spike-RBD (-45 REU) (Figure 3e). The decrease in calculated interface energy for the cryo-EM based models of the complex CVD432/WT-Spike-RBD as compared to that of the complex CVD293/WT-Spike-RBD is surprising for three reasons. First, S34 can adopt several different conformations (Figure 3d). Secondly, S34 makes weaker energetic contributions to the interface energy in all of the cryo-EM based models of the complex CVD432/WT-Spike-RBD than I34 in the complex CVD293/WT-Spike-RBD, regardless of the serine rotamer (Figure S6a). Third, S34 in CVD432 also has a higher average per-residue energy than I34 in CVD293 (Figure S6b). It is possible that a serine was enriched at position 34 in our directed evolution campaign using error-prone PCR because of the isoleucine parental codon, from which single base mutations could only lead to large hydrophobic amino acids, serine, threonine, or asparagine. Of these possibilities, serine is the smallest amino acid, and might have been favored simply to allow the other ACE2 residues to maintain favorable interactions with the RBD.

It appears that no individual residue at the interface is fully responsible for the improved interface energy in CVD432 cryo-EM based models over the CV293 design; rather, this improvement is the sum of several small improvements among many residues at the interface (Figure S6a). Interestingly, CVD432 contains an N33D mutation away from the interface which may be responsible for such overall stabilization of the interface. In all our models, N33D mutation breaks a hydrogen bond with the Q96 side-chain within ACE2 yielding a subtle shift of the neighboring residues towards the RBD (Figure S7b). This is correlated with subtly lower interface energies for these residues and lower total energy for D33 (Figure S6b, c). The N33D mutation was also observed to improve the RBD binding interaction in another ACE2 engineering study (Chan et al., 2020) and a computational analysis of ACE2 mutations (Chowdhury et al., 2020). Additionally, the interface helix may be stabilized by the main chain-side chain hydrogen bond between the S34 hydroxyl group and the F31 backbone carbonyl group in ACE2 (Figure 3d) that is not appropriately scored by Rosetta. Altogether, the improved SARS-CoV-2 neutralization by CVD432 compared to CVD293 is driven by mutations at the interface that improve binding in addition to stability-enhancing mutations outside the interface. As such, both protein stability and interface preorganization contribute to the overall stability of the receptor trap-RBD complex.

### ACE2 receptor traps bind Omicron-RBD with increased affinity and neutralize the SARS-CoV-2 VOC

Several of the 37 Omicron Spike mutations have been observed in other SARS-CoV-2 variants. For example, previous reports suggest that the N501Y mutation increases WT-ACE2 binding affinity while the K417N mutation decreases the WT-ACE2 binding affinity in some SARS-CoV-2 variants (Barton et al., 2021; Laffeber et al., 2021; Liu et al., 2021; Mannar et al., 2021; Tian et al., 2021; Zhu et al., 2021). Fourteen of the 37 mutations have not been reported previously in other variants but together were reported to improve the binding affinity ∼2-3 fold between Omicron-RBD and WT-ACE2 (Cameroni et al., 2022; Mannar et al., 2022; McCallum et al., 2022). For example, while the Y505H mutation leads to loss in hydrogen bond interactions with WT-ACE2 residue E37, there are several compensatory mutations in the Omicron-RBD. These include Q493R and G496S, which result in new hydrogen bonds with WT-ACE2 residues E35 and K353, respectively and several others (Table S1, residue pairs colored green).

Our ACE2 receptor traps have >170-fold improved binding affinity for monomeric WT-Spike-RBD compared to the binding affinity of WT-ACE2 for WT-Spike-RBD (Glasgow et al., 2020). To evaluate if our ACE2 receptor traps would bind Omicron-RBD, we first generated models of Omicron-RBD (330-530)/CVD293 (18-614) and Omicron-RBD (330-530)/CVD432 (18-614) by superimposing and replacing WT-Spike-RBD in our respective cryo-EM local refinement of the complexes with Omicron-RBD from the recently determined cryo-EM structure of Omicron-Spike (330-530)/WT-ACE2 (19-613) (PDB ID: 7T9L, EMD-25761) (Figure 4a,b, panels on the right); figure S8) and minimizing the complexes. Next, we calculated the Rosetta interface energies of these models (Figure 4a,b; panels on the left). Although we predict an energy penalty for engineered interface interaction residue pairs going from R493/E35 in Omicron-RBD/WT-ACE2 to R493/Q35 in Omicron-RBD/CVD293 or Omicron-RBD/CVD432 (in red in Figure 4a-b; panels on the left), overall, we also predict improved binding affinities between the Omicron-RBD and the engineered ACE2 receptor traps. Total interface energy for the interface residue pairs contributing to the affinity between CVD293 or CVD432 to Omicron-RBD was calculated to be - 10.77 REU and -8.5 REU, respectively. For reference, the total interface energy for the interface residue pairs contributing to the affinity between WT-ACE2 and Omicron-RBD was calculated to be -4.99 REU. Potential contributions from interaction residue pairs L455/F31, Y489/F31 and F31/R493 (in yellow in Figure 4a-b; panels on the left) and other residue pair interactions between residue I34 of CVD293 or S34 of CVD432 and L455 of Omicron-RBD may be improving the interface affinity (in blue in Figure 4a-b; panels on the left).

**Figure 4.**
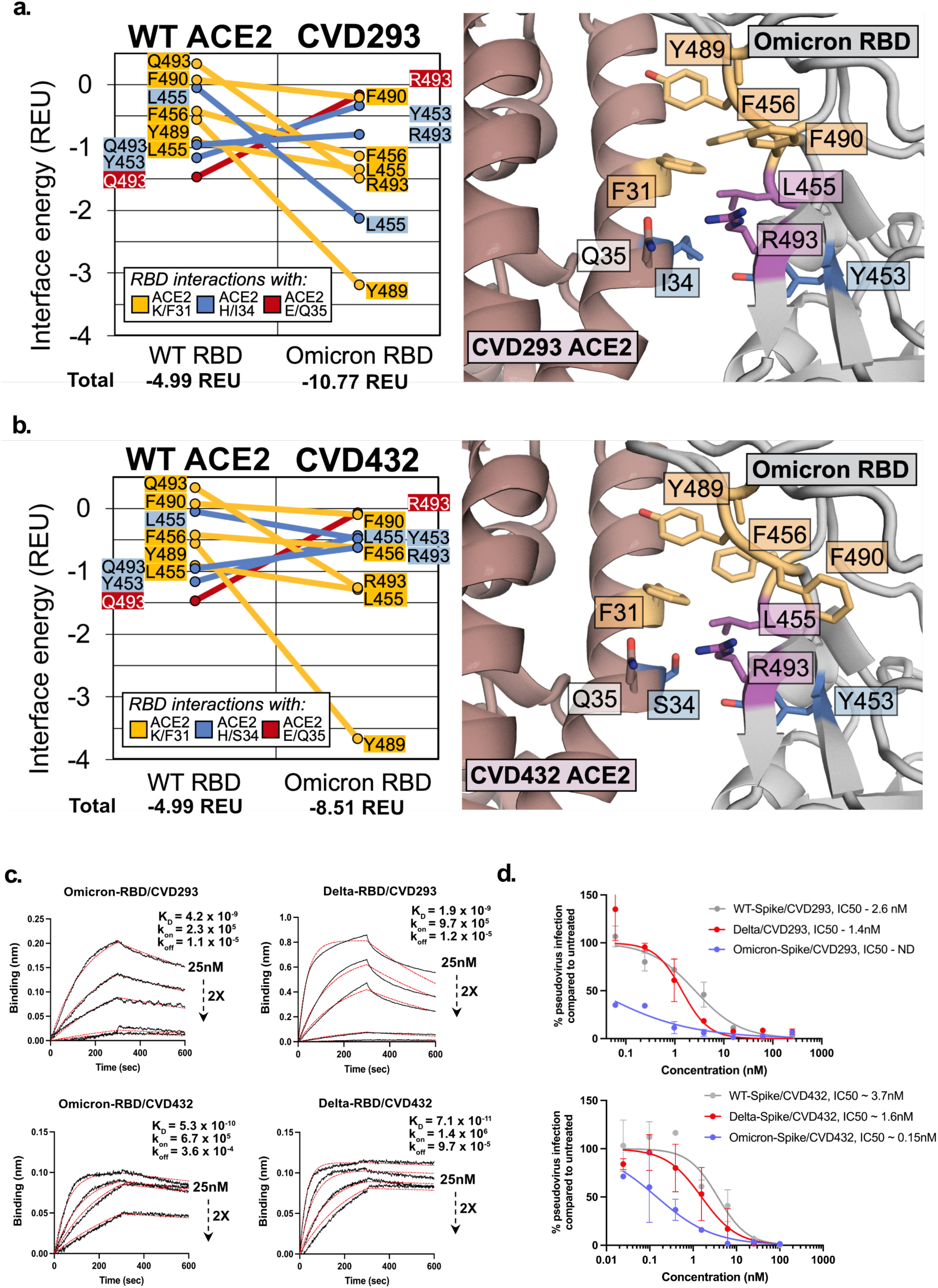
Binding to Omicron- and Delta-RBD and neutralization of Omicron- and Delta-SARS-CoV-2 VOCs by CVD293 and CVD432. **a-b.** Left panel - Predictions based on Rosetta interface energy calculations suggest that Omicron-RBD binds CVD293 and CVD432 with high affinity. Residue pair interactions of RBD residues with CVD293/CVD432 residue F31 (yellow), with residue I34/S34 (blue) and with residue Q35 (red) are shown. Right panel - Zoomed in view of the interface of the models Omicron-RBD/CVD293 and Omicron-RBD/CVD432. Wheat-colored residues indicate RBD interactions ACE2 K/F31. Blue residues indicate RBD interactions with ACE2 H/I/S34. Magenta residues indicate RBD interactions with more than one engineered ACE2 residue **c.** Biolayer interferometry measurements for CVD293 or CVD432 interactions with Omicron- or Delta-RBD. **d.** CVD293 and CVD432 potently neutralize vesicular stomatitis virus (VSV) pseudotyped with SARS-CoV2 Omicron- and Delta-Spike. Error bars represent standard deviation over all technical replicates from two biological replicates.

To test whether improved predicted interface energies correspond to increased apparent binding affinities, we assayed the binding affinity of CVD293 and CVD432 for Omicron-RBD by performing biolayer interferometry (BLI). The BLI-determined dissociation constant (K_D_) between Omicron-RBD/CVD293 (K_D_ = 4.2 nM) or Omicron-RBD/CVD432 (K_D_ = 0.53 nM) were measured to be 10- and 100-fold lower than that for Omicron-RBD/WT-ACE2(18-740)-Fc-fusion, respectively (Figure 4c, Figure S9, Table S2). The hydrophobic interactions specific to Omicron-RBD/ACE2 receptor trap complexes, along with the several compensatory mutations in Omicron-RBD/ACE2 interface that are also maintained with the ACE2 receptor traps, likely result in the BLI-measured improved affinity. Interestingly, CVD293 and CVD432 showed similar (K_D_ = 1.9 nM) or about 100-fold (K_D_ = 0.071 nM) lower K_D_ for Delta-RBD, respectively.

Finally, to determine whether the improved in vitro binding affinity also leads to higher potency in viral neutralization, we tested the neutralization of Omicron (B.1.1.529) and Delta (B.1.617.2) SARS-CoV-2 variants by our ACE2 receptor traps in pseudoviruses bearing these variants of interest generated using recombinant vesicular stomatitis virus (VSV) expressing green fluorescent protein (GFP) in place of the VSV glycoprotein (rVSV ΔG-GFP). We compared the neutralization of Delta and Omicron pseudoviruses to a control Spike-WT pseudovirus with a D614G mutation. The pseudovirus neutralization assays demonstrated that both CVD293 and CVD432 neutralize Delta (IC50 = 1.4 nM, 1.6 nM, respectively) and Omicron (IC50 = ND, 0.15nM, respectively) pseudoviruses, with IC50 values improved between 2-20-fold over Spike-WT (IC50 = 2.6 nM, 3.7 nM, respectively) pseudovirus (Figure 4d). Taken together, the results from the BLI and pseudovirus neutralization assays suggest that our engineered ACE2 traps, although computationally designed and affinity improved against WT-Spike-RBD, can still bind the rapidly evolving SARS-CoV-2 variants with high affinity and potently block virus entry into cells.

## Discussion

Over the course of the current pandemic, several neutralizing monoclonal antibodies (mAbs) have been identified and some have been evaluated clinically as therapeutic candidates against SARS-CoV-2 infection (Barnes et al., 2020a; Baum et al., 2020; Cohen et al., 2021; Dong et al., 2021; Gottlieb et al., 2021; Hansen et al., 2020; Jones et al., 2021; Shi et al., 2020; Zost et al., 2020). These mAbs are broadly categorized based on their ability to bind RBD in the “up” or “down” conformations on fl-Spike, engaging epitopes that can or cannot block ACE2 receptor binding to the RBD (Barnes et al., 2020b, 2020a; Liu et al., 2020; Shi et al., 2020; Tortorici et al., 2020). Several of the mAbs that bind the ACE2 recognition site (also called the receptor binding motif, RBM) within the RBD lost in-vitro neutralization activity against the Omicron-VOC (Cameroni et al., 2022; Mannar et al., 2022; Park et al., 2022). Interestingly, only the S2K146 mAb that binds SARS-CoV-2, SARS-CoV and other sarbecoviruses through ACE2 molecular mimicry retained neutralization activity against the Omicron-VOC (Cameroni et al., 2022; Park et al., 2022). This suggests that ACE2 specific binding epitope residues have a higher barrier for emergence of escape mutants.

We (Glasgow et al., 2020) and others (Chan et al., 2020; Higuchi et al., 2021; Lei et al., 2020) have explored “ACE2 decoy receptors” or “ACE2 receptor traps” that bind and block the RBD as an alternative approach for SARS-CoV-2 virus neutralization. Our ACE2 receptor traps were computationally designed, and affinity matured against the WT-Spike-RBD. In this study, we first determined the cryo-EM structures of the ACE2 receptor traps with the WT-fl-Spike to validate the mechanism of action for our computationally designed trap and to determine the molecular basis for the additional binding affinity improvement by the affinity-matured trap. Although the cryo-EM structures were informative, the limited resolution of ACE2/Spike-RBD interface residues prompted us to explore a multi-model cryo-EM structural workflow to circumvent the limited resolution and leverage model ensembles for interface energy calculations.

From the cryo-EM structures and the derived multi-model workflow, we learned key lessons for designing strong binders: that distributed binding interactions across a protein-protein interface are more effective as compared to reliance on one or two important cross-interface interactions; and that it is important to prioritize the stability of all proteins individually in addition to the protein complex. Importantly, such stability may come from substitutions away from directly interacting residues, like N33D mutation in CVD432 that yields a lower energy conformation for that residue allowing for lower interface energies of surrounding residues. The data suggest that the stability and pre-organization of each protein at the interface is as important as the cross-interface interactions in the overall stability of the complex.

Recently, two engineered ACE2 decoy receptors have been reported to broadly neutralize SARS-CoV-2 variants including the Omicron-VOC (Ikemura et al., 2021; Zhang et al., 2022). These engineered decoy receptors neutralized the Omicron-VOC with IC50s comparable or even better than Omicron-VOC neutralizing mAbs such as VIR-7831. Apeiron’s APN01, a wild-type ACE2 soluble extracellular domain, has also shown promising results in early-phase clinical trials and retains the ability to neutralize multiple variants of concern (Wirnsberger et al., 2021). Furthermore, in a laboratory simulation of viral mutation under neutralizing selective pressure, another engineered ACE2 decoy receptor, 3n39v2 retained its neutralizing capacity over several passage cycles (Higuchi et al., 2021). Thus, soluble engineered ACE2 receptors have therapeutic value against SARS-CoV-2 variants and may continue to be relevant as this virus evolves further.

Application of the multi-model workflow increased our confidence in the atomic positions of computationally designed amino acids at the ACE2-Spike interface. This laid the groundwork for further improvements in the receptor trap design and allowed us to model interactions of the receptor traps with the Omicron-RBD. We experimentally verified the Omicron-RBD binding interactions with the receptor traps using BLI and pseudovirus neutralization assays, demonstrating that our ACE2 receptor traps designed for neutralization of WT-fl-Spike from SARS-CoV-2 remain robust to dozens of mutations in the VOCs. This is both surprising and exciting since our computational design and affinity maturation optimized the binding interface of ACE2 to selectively bind the targeted antigen, WT-Spike-RBD. These results are also in contrast to pan-specific antibodies that are affinity matured to be highly epitope- or antigen-specific binders (Koerber et al., 2013; Mou et al., 2018; Zhou et al., 2020b). Perhaps, the particular hydrophobic mutations at the binding interface of our ACE2 receptor traps make them more adaptable to Spike-RBD mutations. Alternatively, as the virus evolves, its affinity for its entry receptor increases and fortuitously also to our ACE2 receptor traps.

In addition to the RBM antigenic site, other mAbs with antigenic sites outside the RBM such as Sotrovimab, S309, S2X259 and S2H97 also retained neutralization activity against Omicron-VOC (Cameroni et al., 2022; Mannar et al., 2022; McCallum et al., 2022). A bifunctional antibody format called ReconnAbs (receptor-blocking conserved non-neutralizing antibodies) was recently shown to convert non-neutralizing antibodies to potent neutralizers of SARS-CoV-2 VOCs by linking the “WT-ACE2 receptor” to a bispecific antibody targeting two non-overlapping conserved epitopes (Weidenbacher et al., 2022). We envision future versions of our ACE2-receptor trap binders to be knob-in-hole bispecifics and other Fc-fusion formats with one engineered ACE2 arm and other arm(s) as mAbs with antigenic sites outside the RBM, other non-neutralizing mAbs or Vh domains.

Overall, this study exemplifies how technical advances in cryo-EM and computational protein design methods can be combined towards improving the *design-build-test* cycle for engineering potent biotherapeutics, even for difficult targets such as the ACE2 complex with the SARS-CoV-2 Spike protein. Furthermore, this workflow can be generalized for solving the cryo-EM structures of other protein complexes and improving computational protein design protocols.

## Acknowledgements

We thank members of the Wells Lab, particularly those working on COVID-19 projects, for their efforts and contributions. J.A.W. is supported by generous grants from NCI (R35 GM122451-01); the Chan-Zuckerberg Biohub, UCSF Program for Breakthrough Biomedical Research (PBBR); Fast Grants from Emergent Ventures at the Mercatus Center, George Mason University (#2154); and funding from The Harrington Discovery Institute (GA33116). A.G is supported by a grant from NIH (K99GM135529). K.A.V. is supported by Fast Grants from Emergent Ventures at the Mercatus Center, George Mason University and by QBI Independent Research Fellowship.

The structural biology portion of this work was performed by the QCRG (Quantitative Biosciences Institute Coronavirus Research Group) Structural Biology Consortium. Listed below are the contributing members of the consortium listed by teams in order of team relevance to the published work. Within each team the team leads are italicized (responsible for organization of each team, and for the experimental design utilized within each team), then the rest of team members are listed alphabetically.

## QCRG Structural Biology Consortium

### CryoEM grid freezing/collection team

*Axel F. Brilot, Gregory E. Merz, Alexandrea N. Rizo,* Caleigh M. Azumaya, Julian R. Braxton, Meghna Gupta, Fei Li, Kyle E. Lopez, Arthur Melo, Frank Moss, Joana Paulino, Thomas H. Pospiech Jr., Sergei Pourmal, Cristina Puchades, Amber M. Smith, Ming Sun, Paul V. Thomas, Feng Wang & Zanlin Yu

### CryoEM data processing team

*Axel F. Brilot, Gregory E. Merz, Alexandrea N. Rizo, Soumya G. Remesh,* Daniel Asarnow, Julian R. Braxton, Melody G. Campbell, Cynthia M. Chio, Un Seng Chio, Miles Sasha Dickinson, Devan Diwanji, Bryan Faust, Meghna Gupta, Nick Hoppe, Mingliang Jin, Fei Li, Junrui Li, Yanxin Liu, Henry C. Nguyen, Joana Paulino, Thomas H. Pospiech Jr., Sergei Pourmal, Smriti Sangwan, Raphael Trenker, Donovan Trinidad, Eric Tse, Kaihua Zhang & Fengbo Zhou

### Mammalian cell expression team

*Evelyn Hernandez, Devan Diwanji, Amber M. Smith,* Caleigh M. Azumaya, Christian Billesboelle, Alisa Bowen, Melody G. Campbell, Nick Hoppe, Yen-Li Li, Edmond Linossi, Jocelyne Lopez, Phuong Nguyen, Carlos Nowotny, Quynh Mai, Hevatib Mehmood, Michael Paul, Cristina Puchades, Mali Safari, Smriti Sangwan, Kaitlin Schaefer, Raphael Trenker, Tsz Kin Martin Tsui, Natalie Whitis & Jianhua Zhao

### Protein purification team

*Michelle Moritz, Sergei Pourmal,* Daniel Asarnow, Caleigh M. Azumaya, Cynthia M. Chio, Bryan Faust, Meghna Gupta, Raghav Kalia, Kate Kim, Tristan W. Owens, Joana Paulino, Komal Pawar, Jessica K. Peters, Kaitlin Schaefer & Tsz Kin Martin Tsui

### Crystallography team

Nadia Herrera, Huong T. Kratochvil, Ursula Schulze-Gahmen, Iris D. Young, Justin Biel, Ishan Deshpande & Xi Liu

### Bacterial expression team

*Amy Diallo, Meghna Gupta,* Jen Chen, Loan Doan, Sebastian Flores, Mingliang Jin, Huong T. Kratochvil, Victor L. Lam, Yang Li, Megan Lo, Gregory E. Merz, Joana Paulino, Aye C. Thwin, Erron W. Titus, Zanlin Yu, Fengbo Zhou & Yang Zhang

### Infrastructure team

David Bulkley, Arceli Joves, Almarie Joves, Liam McKay, Mariano Tabios & Eric Tse

### Leadership team

David A. Agard, Yifan Cheng, James S. Fraser, Adam Frost, Natalia Jura, Tanja Kortemme, Nevan J. Krogan, Aashish Manglik, Oren S. Rosenberg, Daniel R. Southworth, Robert M. Stroud & Kliment A. Verba

## Author Contributions

Conceptualization, K.A.V, A.G., S.G.R, T.K., K.K.L., J.A.W.; methodology (cryo-EM data collection), G.E.M, A.F.B., A.N.R., T.H.P.; methodology (cryo-EM structure solution), G.E.M, A.F.B., U.S.C., S.G.R.; methodology (cryo-EM based multi-model workflow), S.G.R. and K.A.V.; methodology (Rosetta energy calculations), A.G.; methodology (cloning, protein expression and purification), J.G., S.G.R., I.L., C.Q.L., Y.Z., E.H., D.D., S.P., A.M.S., J.L., K.I.P.; methodology (Biolayer interferometry), I.L.; methodology (pseudovirus generation), M.T.L., J.D.; methodology (pseudoviral assay), S.G.R.; writing—original draft, S.G.R., A.G. K.A.V., T.K., J.A.W., K.K.L.; visualization, S.G.R., A.G., K.A.V.; supervision; K.A.V., A.G., K.K.L., J.A.W.; funding acquisition, K.A.V., A.G., J.A.W

## Materials and Methods

### Cloning, Expression and Purification of Protein Constructs

WT-fl-Spike plasmid was a generous gift from the Pak lab (Chan Zuckerberg Initiative Biohub) and Krammer lab (Icahn School of Medicine at Mount Sinai). WT-fl-Spike construct has an N-terminal spike protein signal peptide, a trimerization domain and C-terminal 6XHis-tag. WT, Delta-, Omicron-SARS-CoV-2 spike RBD and ACE2 variants (CVD293) were cloned into a pFUSE-based vector with Zeocin antibiotic resistance marker for mammalian expression using the Gibson method. The SARS-CoV-2 spike RBD construct has an IL2 secretion signal followed by the gene of interest, a Gly-Ser linker, a TEV protease cut site, a Gly-Ser linker, an 8X His-tag and an Avitag. CVD293 was cloned into a similar construct with N-terminal IL2 secretion signal followed by ACE2 with the relevant mutations (K31F, H34I, E35Q), a Gly-Ser linker, a TEV protease cut site, a Gly-Ser linker, a human IgG1 hinge and Fc, and AviTag. Construct CVD432 was purchased from Twist Bioscience (www.twistbioscience.com). The construct has an Ampicillin resistance marker, an N-terminal ACE2 secretion signal followed by ACE2 with the relevant CVD313 mutations (K31F, N33D, H34S, E35Q), a short Gly-Ser linker, a human IgG1 hinge and Fc. Also, the H345L mutation in CVD313 was reverted to wildtype residue, histidine.

WT-fl-Spike protein was purified based on the previously described protocol (Weinberg et al., 2021) 30ml of ExpiCHO-S cells at 6M cells/mL were transfected with 1ug/mL of the spike WT using the ExpiCHO^TM^ Expression System Kit (Gibco, catalog # A29133) following the manufacturer’s Standard protocol. Briefly, the transfected cells were shaken at 37°C and 8% CO_2_ and 18 hours post-transfection, the cultures were supplemented with ExpiCHO^TM^ Feed and ExpiFectamine^TM^ CHO Enhancer and continued to be shaken at 37°C and 8% CO_2_ for up to 10 days. The supernatant with secreted spike protein was collected by centrifugation of the culture at 4000xG for 15 mins. The media was filtered using a 0.42µM filter and was transferred to a 50 mL centrifuge tube. The sample was then mixed with 2 mL washed Ni-Excel resin (Millipore Sigma, catalog # GE17371201) pre-equilibrated with 10 mM Tris/HCl, pH 8.0, 200 mM NaCl and placed on a rocker for 1 hour at room temperature. After binding, the sample was washed with 25 bed volumes of Wash buffer (10 mM Tris/HCl, pH 8.0, 200 mM NaCl, 10 mM imidazole), and eluted with 7 bed volumes of elution buffer (10 mM Tris pH 8.0, 200 mM NaCl, 500 mM imidazole) into separate 2 mL centrifuge tubes. The eluent was concentrated, filtered, and a final SEC polishing step was performed on Superose^®^ 6 10/300 Increase (Millipore Sigma, catalog # GE17-5172-01) pre-equilibrated in 10 mM HEPES pH 8.0, 200 mM NaCl at 4°C.

WT, Delta-, Omicron-SARS-CoV-2 spike RBD and ACE2 variants (CVD293 and CVD432) were expressed in high density Expi293F^TM^ cells in Expi293^TM^ expression media following manufacturer’s protocol (Expi293 Expression System, ThermoFisher Scientific). Briefly, 3x10^6^ cell/ml at >95% viability in 25ml media were transfected with ∼30ug of DNA using ExpiFectamine^TM^ transfection reagent and Opti-MEM^TM^ I medium. The transfected cells were incubated at 37 °C, 8% CO_2_ on an orbital shaker, supplemented with ExpiFectamine^TM^ 293 Transfection Enhancer 1 and ExpiFectamine^TM^ 293 Transfection Enhancer 2 about 18-22 hours post-transfection and continued to be incubated at 37 °C, 8% CO_2_ on an orbital shaker for additional 4-5 days. Cells were harvested 4-5 days post-transfection, centrifuged at 3000 x g for 15 minutes, supernatant collected and filtered through 0.22µM syringe filter. Proteins were neutralized with 10X phosphate buffered saline (PBS, 0.01 M phosphate buffer, 0.0027 M KCl and 0.137 M NaCl, Millipore Sigma P4417-100TAB), pH 7.4 to a final concentration of 2.5x PBS (342.5 mM NaCl, 6.75 mM KCl and 29.75 mM phosphates). SARS-CoV-2 RBDs were purified using cobalt-based immobilized metal affinity chromatography followed by buffer exchange into 1X PBS using a Superdex 200 Increase 10/300 GL column (Cytiva). Fc-fused ACE2 proteins were purified on HiTrap Protein A column (GE Healthcare) and eluted with 50 mM Tris pH 7.2, 4 M MgCl_2_ and buffer exchanged into 1X PBS. The protein concentrations were estimated based on the protein absorbance at 280 nm with a spectrophotometer (Nanodrop One, Thermo). All the proteins were >95% pure as determined by SDS-Page gel electrophoresis. The proteins were aliquoted, flash frozen, and stored in -80°C.

### Cryo-electron Microscopy Sample Preparation and Data Collection

Purified WT-fl-Spike at 2 μM was mixed with either CVD293 (at 2.5 μM) or CVD432 (at 3 μM) just prior to plunge freezing. 300 mesh 1.2/1.3R UltrAufoil grids were glow discharged at 15 mA for 30 seconds. Vitrification was done using FEI Vitrobot Mark IV (ThermoFisher) set up at 4°C and 100% humidity. 4 μl complex sample was applied to the grids and the blotting was performed with a blot force of 0 for 4 s prior to plunge freezing into liquid ethane. For WT-fl-Spike/CVD293 complex, two datasets comprising of 2,058 and 3,636 120-frame super-resolution movies each were collected while for WT-fl-Spike/CVD432 complex, three datasets comprising of 4,915, 3,196, and 2,574 120-frame super-resolution movies each were collected. For both complexes movies were acquired in super-resolution mode on a Titan Krios (ThermoFisher) equipped with a K3 camera and a Bioquantum energy filter (Gatan) set to a slit width of 20 eV. Collection was performed with a 3x3 image shift at a calibrated magnification of 105, 000 x corresponding to a super resolution pixel size of 0.4265 Å/pix. Collection dose rate was 8 e^-^/physical pixel/second for a total dose of 68 e^-^/Å2. Defocus range was -0.8 to -1.8 μm. Movies were subsequently corrected for drift using MotionCor2 (Zheng et al., 2017) and were Fourier-cropped by a factor of 2 to a final pixel size of 0.834 Å/pixel. Each collection was performed with semi-automated scripts in SerialEM (Mastronarde, 2003, 2005).

### Data Processing -

#### WT-fl-Spike/CVD293 complex 1-RBD-up state (Figure S2)

Initial processing was done in cryoSPARC (v2.15.0) (Punjani et al., 2017, 2020). The two datasets with 3636 and 2058 dose-weighted motion corrected micrographs (Zheng et al., 2017) were imported and Patch CTF(M) was performed. Micrographs were curated based on CTF-fit resolution (<4 Å), ice-thickness, and presence of carbon. After manual curation, 3144 and 1575 micrographs were selected for further processing. Blob-picker was used to pick 1,673,305 and 831,773 particles and extraction was done with a box size of 580 px downsized to 480 px for each dataset. 2D-classification was done into 150 classes and good classes were selected with a total of 401,671 particles combined from both datasets. Multiple rounds of heterogeneous, homogenous, and non-uniform refinements in cryoSPARC and focused classification in cisTEM (Grant et al., 2018) resulted in a 3.77 Å 3D-reconstruction of WT-fl-Spike/CVD293 with 61,033 particles. Particle subtraction/local refinement was performed on this final stack of particles focusing on the CVD293/RBD to obtain a 3.50 Å 3D reconstruction of WT-Spike-RBD (residues 331-541)/CVD293 (residues 18-614).

#### WT-fl-Spike/CVD432 complex 1-RBD-up state (Figure S3)

Initial processing was done in cryoSPARC (v2.15.0). The three datasets with 4915, 3916 and 2574 dose-weighted motion corrected micrographs were imported and Patch CTF(M) was performed. Micrographs were curated based on CTF-fit resolution (<4 Å), ice-thickness, and presence of carbon. After manual curation, 3864, 2238 and 2148 micrographs were selected for further processing. Blob-picker was used to pick 1,259,441 particles from Dataset 1 while template-based particle picker (260 Å diameter) was used to pick 782,620 and 752,648 particles for Dataset 2 and Dataset 3, respectively. Extraction was done with a box size of 512 px for Dataset 1 which was re-extracted with box size of 600 px after a round of ab-initio refinement and finally downsampled to 400 px. For Datasets 2 and 3, extraction was done at box size 600 px and then downsampled to 400 px. 2D-classification was done with 150 classes for each dataset and good classes of WT-fl-Spike/CVD432 in 1-RBD-up state were selected with a total of 601,624 particles combined from the three datasets. Multiple rounds of heterogeneous, non-uniform refinements and homogenous refinements in cryoSPARC and focused classification in cisTEM resulted in a 3.5 Å 3D-reconstruction of WT-fl-Spike/CVD432 1-RBD-up state with 97,082 particles. Particle subtraction/local refinement was performed on this final stack of particles focusing on the CVD432/RBD to obtain a 3.36 Å 3D reconstruction of WT-Spike-RBD (residues 331-541)/CVD432 (residues 18-614).

#### WT-fl-Spike/CVD432 complex 2-RBD-up state (Figure S4)

We processed the WT-fl-Spike/CVD432 complex 2-RBD-up state independent of the 1-RBD-up state. Initial processing was done in cryoSPARC (v2.15.0). The three datasets with 4915, 3916 and 2574 dose-weighted motion corrected micrographs were imported and Patch CTF(M) was performed. Micrographs were curated based on CTF-fit resolution (<4 Å), ice-thickness, and presence of carbon. After manual curation, 3833, 2253 and 2175 micrographs were selected for further processing. We first used blob-picker on Dataset 1 to pick 1,597,143 and after 2D classification, 19 good classes were selected. The 19 classes from Data set 1 were used for template-based particle picker (220 Å diameter) to pick 1,499,401 and 1,019,564 and 1,328,287 particles for Dataset 1, Dataset 2 and Dataset 3, respectively. Extraction was done at box size 600 px for all datasets. 2D-classification was done into 150 classes for each dataset and good looking classes with WT-fl-Spike/CVD432 in 2-RBD-up state were selected with a total of 113,836 particles combined from the three datasets. Multiple rounds of heterogeneous, non-uniform refinements and homogenous refinements in cryoSPARC and focused classification in cisTEM resulted in a 2.97 Å 3D-reconstruction of WT-fl-Spike/CVD432 2-up state with 97,374 particles. Each WT-Spike-RBD/CVD432 interface in the 2-RBD-up state was at an overall low resolution and particle subtraction/local refinement was not performed.

### Model Building and Refinement -

#### Low resolution rigid body fitting

1-RBD-up state WT-fl-Spike/WT-ACE2 model with PDB ID - 7DX5, was used as the initial model for rigid body fitting into cryo-EM density map of 1-RBD-up state of WT-fl-Spike/CVD293 or 1-RBD-up state of WT-fl-Spike/CVD432 in CHIMERA (Pettersen et al., 2004). For 2-RBD-up state of WT-fl-Spike/CVD432, we generated a model by rigid body fitting PDB ID - 7DX5 into the density map and simultaneously rigid body fitting WT-Spike-RBD/ACE2 (18-614) complex with PDB ID - 6M0J into the second RBD-up region of the map in CHIMERA. The final model was fit using FastRelax in torsion space in Rosetta into the WT-fl-Spike/WT-ACE2 2-RBD-up state cryo-EM map. Rosetta into the Overall, the fits agree with previously reported 1-RBD-up states for WT-fl-Spike/ACE2 (18-614) complexes.

### High resolution “final consensus” model building

High resolution map of WT-Spike-RBD/CVD293 (18-614) with mutations K31F, H34I, E35Q or WT-Spike-RBD/CVD432 (18-614) with mutations K31F, N33D, H34S, E35Q were generated from WT-fl-Spike/CVD293 map or WT-fl-Spike/CVD432 map, respectively, in cryoSPARC (v2.15.0) by Particle subtraction/Local refinement. PDB ID: 6M0J was used for initial rigid body fit into the WT-Spike-RBD/CVD293 (18-614) map. This model was then refined against the respective maps in Phenix Real Space Refine (Liebschner et al., 2019) with the corrected sequence input for CVD293 or CVD432. This was followed by a FastRelax in torsion space with Rosetta (2020.08 release) (Wang et al., 2016). Model for each complex was manually examined and corrected using COOT 0.9 (Emsley et al., 2010) and ISOLDE 1.0 (Croll, 2018). The B-factors were assigned using a Rosetta B-factor fitting mover. Local resolution was determined by running the ResMap program (Kucukelbir et al., 2014). Directional FSC curves were determined by submitting the associated files to the 3DFSC server (Tan et al., 2017). Q-scores were calculated using the Q-score plugin for USCF Chimera (Pintilie et al., 2020).

### Multi-model Pipeline

The Cryo-EM consensus model of WT-Spike-RBD/CVD293 (18-614) or WT-Spike-RBD/CVD432 (18-614) was used as the starting model for the pipeline. We applied the Rosetta Iterative Rebuild protocol on 10-amino acid overlapping stretches of the interface helix for each consensus model. The parameters used were defined in an XML file (Parameters.xml) found in SI Appendix: Supplemental computational methods: Commands and input files section. For each 10-amino acid stretch of the interface helix we generated 2000 independent models totaling 8000 models for the entire interface helix from ACE2 residue 21 to residue 52. Total of 200 models were then selected based on the total Rosetta energy score followed by the top 20 that best fit to the map destiny. A total of 80 top selected models for each complex were then used for further analyses. We calculated the local average per residue average side-chain RMSD in CHIMERA and average interface energy (in REU) for the residues of the interface helix.

### Energy calculations

Total and individual pairwise interface energies were calculated for all design models and cryo-EM atomic models using the Rosetta interface energy application. Per-residue energies were calculated using the Rosetta per-residue energy application, and total energies were calculated using the Rosetta scoring application in Rosetta (2021.48.post.dev+8.master.77491fa20be77491fa20be83588cfc37ab422ba5b95eca128ebgit@ github.com:RosettaCommons/main.git 2021-12-02T08:56:13) (Alford et al., 2017; Leman et al., 2020) . For the cryo-EM atomic models of CVD293 and CVD432, all scores were averaged and standard deviations were calculated. The command lines and code are available in the *SI Appendix: Supplemental computational methods: Commands and input files section*.

### Design models of the ACE2 domain of CVD432

Design models of the ACE2 domain of CVD432 were generated using the RosettaScripts framework (Glasgow et al., 2020). Beginning with the atomic models of CVD293 solved by cryo-EM using the multi-model pipeline, we mutated N33 and I34 to aspartic acid and serine, respectively, in each of the 80 WT-Spike-RBD/CVD293 cryo-EM based models and minimized the complexes. Total energies and interface energies were calculated as described in “Energy calculations.” The average interface energies and standard deviation is shown as CVD293 cryo-EM based models in Figure 3e. The code is available in the *SI Appendix: Supplemental computational methods: Commands and input files section*.

### Determination of binding affinity using biolayer interferometry (BLI)

Affinity measurements were carried out at room temperature using an Octet RED384 system. In our BLI experiments, ACE2 Fc-fusions are tethered to either Streptavidin biosensors (WT-ACE2, CVD293) (Sartorius®, Item no.: 18-5019) or ProtA biosensors (Sartorius®, Item no.: 18-5010) and WT, Delta-, Omicron-SARS-CoV-2 spike RBD are present as the analyte in solution in a 384-well microplate. The biosensors were washed in phosphate buffered saline (1X PBS) with 0.05% Tween-20 at pH 7.4 and 0.2% bovine serum albumin (BSA) (1X PBSTB) for 200 seconds. Antigens WT-ACE2, CVD293 were diluted to 10 nM in 1X PBSTB containing 10 µM biotin (1X PBSTBB) as blocking agent while antigen CVD432 was diluted 10nM in 1X PBSTB. The antigens were then loaded to the respective biosensors for 300 seconds. Following loading, a baseline was established by washing the WT-ACE2 or CVD293 bound Streptavidin biosensors in 1X PBSTBB and CVD432 bound ProtA biosensors in 1X PBSTB for 200 seconds. WT, Delta-, Omicron-SARS-CoV-2 spike RBD were then allowed to associate with the antigen at concentrations ranging from 0 to 25 nM of Spike for about 600 seconds and then returned to the respective washing well to follow dissociation for about 900 seconds. Raw data were fit in Octet Data Analysis HT software version 10.0 using curve-fitting kinetic analysis with global fitting and assuming 1:1 non-cooperative binding kinetics. All fits to BLI data had R^2^ (goodness of fit) > 0.90.

### Pseudovirus Production

SARS-CoV-2 pseudoviruses bearing spike proteins of variants of interest were generated as previously described (Hoffmann et al., 2020; Laurie et al., 2021, 2022) using a recombinant vesicular stomatitis virus (VSV) expressing green fluorescent protein (GFP) in place of the VSV glycoprotein (rVSV Δ G-GFP). B.1 (WT, 1 spike mutation (D614G)), B.1.617.2/delta (9-10 spike mutations), and B.1.1.529/ omicron (37 spike mutations) were cloned in a cytomegalovirus enhancer-driven expression vector and used to produce SARS-CoV-2 spike pseudoviruses. The mutations for each variant are listed as follows - B.1/WT (D614G), B.1.617.2/Delta (T19R, T95I, G142D, Δ157-158, L452R, T478K, P681R, D614G, D950N), B.1.1.529/Omicron (A67V, ΔH69, ΔV70, T95I, ΔG142, ΔV143, ΔY144, Y145D, ΔN211, L212I, G339D, S371L, S373P, S375F, K417N, N440K, G446S, S477N, T478K, E484A, Q493R, G496S, Q498R, N501Y, Y505H, T547K, D614G, H655Y, N679K, P681H, N764K, D796Y, N856K, Q954H, N969K, L981F). Pseudoviruses were titered on Huh7.5.1 cells overexpressing ACE2 and TMPRSS2 (gift of Andreas Puschnik) using GFP expression to measure the concentration of focus forming units (ffu).

### Pseudovirus Neutralization Assay

Pseudovirus Neutralization Assay was performed as described previously (Laurie et al., 2021, 2022) Huh7.5.1-ACE2-TMPRSS2 cells were seeded in 96-well plates at a density of 7000-8000 cells/well 1 day prior to pseudovirus infection. ACE2 receptor traps were serially diluted into complete culture media (Dulbecco’s Modified Eagle’s Medium with 10% fetal bovine serum, 10mM HEPES, 1X Pen-Strep-Glutamine). Each pseudovirus was diluted to 125 ffu/µL and 30 µL of each pseudovirus was mixed with ∼30 µL of ACE2 receptor traps or media only. Media only with no pseudovirus served as an additional control. These were incubated at 37°C for 1 hour before adding 50 µL of virus:binder incubated mix directly to previously plated cells. Cells inoculated with ACE2 receptor traps/pseudovirus mixtures were incubated at 37°C and 5% CO2 for 24 hours. The cells were then lifted and resuspended using 20 µL of 10X TrypLE Select (Gibco) and GFP signal of infected cells recorded using Beckman CytoFlex Cytometer B4R3V4.

Data were analyzed with FlowJo^TM^ to determine percent GFP-positive pseudovirus transduced cells. GFP-signals from wells with no ACE2 receptor traps (media only with pseudovirus) were used to normalize the data and determine percent neutralization. A 7-8 point dose-response curve was generated in GraphPad Prism and IC50s reported in µg/mL. The reported IC50s are based on two technical and biological replicates for each pseudovirus/ACE2 receptor trap pair.

## Bibliography

Alford, R.F., Leaver-Fay, A., Jeliazkov, J.R., O’Meara, M.J., DiMaio, F.P., Park, H., Shapovalov, M.V., Renfrew, P.D., Mulligan, V.K., Kappel, K., et al. (2017). The Rosetta All-Atom Energy Function for Macromolecular Modeling and Design. J. Chem. Theory Comput. 13, 3031–3048. https://doi.org/10.1021/acs.jctc.7b00125.

Barnes, C.O., West, A.P., Huey-Tubman, K.E., Hoffmann, M.A.G., Sharaf, N.G., Hoffman, P.R., Koranda, N., Gristick, H.B., Gaebler, C., Muecksch, F., et al. (2020a). Structures of Human Antibodies Bound to SARS-CoV-2 Spike Reveal Common Epitopes and Recurrent Features of Antibodies. Cell 182, 828–842.e16. https://doi.org/10.1016/j.cell.2020.06.025.

Barnes, C.O., Jette, C.A., Abernathy, M.E., Dam, K.-M.A., Esswein, S.R., Gristick, H.B., Malyutin, A.G., Sharaf, N.G., Huey-Tubman, K.E., Lee, Y.E., et al. (2020b). SARS-CoV-2 neutralizing antibody structures inform therapeutic strategies. Nature 588, 682–687. https://doi.org/10.1038/s41586-020-2852-1.

Barton, M.I., MacGowan, S.A., Kutuzov, M.A., Dushek, O., Barton, G.J., and van der Merwe, P.A. (2021). Effects of common mutations in the SARS-CoV-2 Spike RBD and its ligand, the human ACE2 receptor on binding affinity and kinetics. ELife 10. https://doi.org/10.7554/eLife.70658.

Baum, A., Fulton, B.O., Wloga, E., Copin, R., Pascal, K.E., Russo, V., Giordano, S., Lanza, K., Negron, N., Ni, M., et al. (2020). Antibody cocktail to SARS-CoV-2 spike protein prevents rapid mutational escape seen with individual antibodies. Science 369, 1014–1018. https://doi.org/10.1126/science.abd0831.

Cameroni, E., Bowen, J.E., Rosen, L.E., Saliba, C., Zepeda, S.K., Culap, K., Pinto, D., VanBlargan, L.A., De Marco, A., di Iulio, J., et al. (2022). Broadly neutralizing antibodies overcome SARS-CoV-2 Omicron antigenic shift. Nature 602, 664–670. https://doi.org/10.1038/s41586-021-04386-2.

Cao, Y., Wang, J., Jian, F., Xiao, T., Song, W., Yisimayi, A., Huang, W., Li, Q., Wang, P., An, R., et al. (2022). Omicron escapes the majority of existing SARS-CoV-2 neutralizing antibodies. Nature 602, 657–663. https://doi.org/10.1038/s41586-021-04385-3.

Cele, S., Jackson, L., Khoury, D.S., Khan, K., Moyo-Gwete, T., Tegally, H., San, J.E., Cromer, D., Scheepers, C., Amoako, D., et al. (2021). SARS-CoV-2 Omicron has extensive but incomplete escape of Pfizer BNT162b2 elicited neutralization and requires ACE2 for infection. MedRxiv https://doi.org/10.1101/2021.12.08.21267417.

Chan, K.K., Dorosky, D., Sharma, P., Abbasi, S.A., Dye, J.M., Kranz, D.M., Herbert, A.S., and Procko, E. (2020). Engineering human ACE2 to optimize binding to the spike protein of SARS coronavirus 2. Science 369, 1261–1265. https://doi.org/10.1126/science.abc0870.

Chowdhury, R., Boorla, V.S., Maranas, C.D. (2020). Computational biophysical characterization of the SARS-CoV-2 spike protein binding with the ACE2 receptor and implications for infectivity. Comput Struct Biotechnol J. 18, 2573–2582.

Cohen, M.S., Nirula, A., Mulligan, M.J., Novak, R.M., Marovich, M., Yen, C., Stemer, A., Mayer, S.M., Wohl, D., Brengle, B., et al. (2021). Effect of Bamlanivimab vs Placebo on Incidence of COVID-19 Among Residents and Staff of Skilled Nursing and Assisted Living Facilities: A Randomized Clinical Trial. JAMA 326, 46–55. https://doi.org/10.1001/jama.2021.8828.

Croll, T.I. (2018). ISOLDE: a physically realistic environment for model building into low-resolution electron-density maps. Acta Crystallogr. D Struct. Biol. 74, 519–530. https://doi.org/10.1107/S2059798318002425.

DiMaio, F., Song, Y., Li, X., Brunner, M.J., Xu, C., Conticello, V., Egelman, E., Marlovits, T., Cheng, Y., and Baker, D. (2015). Atomic-accuracy models from 4.5-Å cryo-electron microscopy data with density-guided iterative local refinement. Nat. Methods 12, 361–365. https://doi.org/10.1038/nmeth.3286.

Dong, J., Zost, S.J., Greaney, A.J., Starr, T.N., Dingens, A.S., Chen, E.C., Chen, R.E., Case, J.B., Sutton, R.E., Gilchuk, P., et al. (2021). Genetic and structural basis for SARS-CoV-2 variant neutralization by a two-antibody cocktail. Nat. Microbiol. 6, 1233–1244. https://doi.org/10.1038/s41564-021-00972-2.

Emsley, P., Lohkamp, B., Scott, W.G., and Cowtan, K. (2010). Features and development of Coot. Acta Crystallogr. D Biol. Crystallogr. 66, 486–501. https://doi.org/10.1107/S0907444910007493.

Glasgow, A., Glasgow, J., Limonta, D., Solomon, P., Lui, I., Zhang, Y., Nix, M.A., Rettko, N.J., Zha, S., Yamin, R., et al. (2020). Engineered ACE2 receptor traps potently neutralize SARS-CoV-2. Proc Natl Acad Sci USA 117, 28046–28055. https://doi.org/10.1073/pnas.2016093117.

Gottlieb, R.L., Nirula, A., Chen, P., Boscia, J., Heller, B., Morris, J., Huhn, G., Cardona, J., Mocherla, B., Stosor, V., et al. (2021). Effect of Bamlanivimab as Monotherapy or in Combination With Etesevimab on Viral Load in Patients With Mild to Moderate COVID-19: A Randomized Clinical Trial. JAMA 325, 632–644. https://doi.org/10.1001/jama.2021.0202.

Grant, T., Rohou, A., and Grigorieff, N. (2018). cisTEM, user-friendly software for single-particle image processing. ELife 7, e35383. https://doi.org/10.7554/eLife.35383.

Hansen, J., Baum, A., Pascal, K.E., Russo, V., Giordano, S., Wloga, E., Fulton, B.O., Yan, Y., Koon, K., Patel, K., et al. (2020). Studies in humanized mice and convalescent humans yield a SARS-CoV-2 antibody cocktail. Science 369, 1010–1014. https://doi.org/10.1126/science.abd0827.

Herzik, M.A., Fraser, J.S., and Lander, G.C. (2019). A Multi-model Approach to Assessing Local and Global Cryo-EM Map Quality. Structure 27, 344–358.e3. https://doi.org/10.1016/j.str.2018.10.003.

Higuchi, Y., Suzuki, T., Arimori, T., Ikemura, N., Mihara, E., Kirita, Y., Ohgitani, E., Mazda, O., Motooka, D., Nakamura, S., et al. (2021). Engineered ACE2 receptor therapy overcomes mutational escape of SARS-CoV-2. Nat. Commun. 12, 3802. https://doi.org/10.1038/s41467-021-24013-y.

Hoffmann, M., Kleine-Weber, H., and Pöhlmann, S. (2020). A Multibasic Cleavage Site in the Spike Protein of SARS-CoV-2 Is Essential for Infection of Human Lung Cells. Mol. Cell 78, 779–784.e5. https://doi.org/10.1016/j.molcel.2020.04.022.

Ikemura, N., Taminishi, S., Inaba, T., Arimori, T., Motooka, D., Katoh, K., Kirita, Y., Higuchi, Y., Li, S., Itoh, Y., et al. (2021). Engineered ACE2 counteracts vaccine-evading SARS-CoV-2 Omicron variant. BioRxiv https://doi.org/10.1101/2021.12.22.473804.

Jones, B.E., Brown-Augsburger, P.L., Corbett, K.S., Westendorf, K., Davies, J., Cujec, T.P., Wiethoff, C.M., Blackbourne, J.L., Heinz, B.A., Foster, D., et al. (2021). The neutralizing antibody, LY-CoV555, protects against SARS-CoV-2 infection in nonhuman primates. Sci. Transl. Med. 13. https://doi.org/10.1126/scitranslmed.abf1906.

Koerber, J.T., Thomsen, N.D., Hannigan, B.T., Degrado, W.F., and Wells, J.A. (2013). Nature-inspired design of motif-specific antibody scaffolds. Nat. Biotechnol. 31, 916–921. https://doi.org/10.1038/nbt.2672.

Kucukelbir, A., Sigworth, F.J., and Tagare, H.D. (2014). Quantifying the local resolution of cryo-EM density maps. Nat. Methods 11, 63–65. https://doi.org/10.1038/nmeth.2727.

Laffeber, C., de Koning, K., Kanaar, R., and Lebbink, J.H.G. (2021). Experimental Evidence for Enhanced Receptor Binding by Rapidly Spreading SARS-CoV-2 Variants. J. Mol. Biol. 433, 167058. https://doi.org/10.1016/j.jmb.2021.167058.

Laurie, M.T., Liu, J., Sunshine, S., Peng, J., Black, D., Mitchell, A.M., Mann, S.A., Pilarowski, G., Zorn, K.C., Rubio, L., et al. (2021). SARS-CoV-2 variant exposures elicit antibody responses with differential cross-neutralization of established and emerging strains including Delta and Omicron. MedRxiv https://doi.org/10.1101/2021.09.08.21263095.

Laurie, M.T., Liu, J., Sunshine, S., Peng, J., Black, D., Mitchell, A.M., Mann, S.A., Pilarowski, G., Zorn, K.C., Rubio, L., et al. (2022). SARS-CoV-2 Variant Exposures Elicit Antibody Responses With Differential Cross-Neutralization of Established and Emerging Strains Including Delta and Omicron. J. Infect. Dis. 225, 1909–1914. https://doi.org/10.1093/infdis/jiab635.

Lei, C., Qian, K., Li, T., Zhang, S., Fu, W., Ding, M., and Hu, S. (2020). Neutralization of SARS-CoV-2 spike pseudotyped virus by recombinant ACE2-Ig. Nat. Commun. 11, 2070. https://doi.org/10.1038/s41467-020-16048-4.

Leman, J.K., Weitzner, B.D., Lewis, S.M., Adolf-Bryfogle, J., Alam, N., Alford, R.F., Aprahamian, M., Baker, D., Barlow, K.A., Barth, P., et al. (2020). Macromolecular modeling and design in Rosetta: recent methods and frameworks. Nat. Methods 17, 665–680. https://doi.org/10.1038/s41592-020-0848-2.

Liebschner, D., Afonine, P.V., Baker, M.L., Bunkóczi, G., Chen, V.B., Croll, T.I., Hintze, B., Hung, L.W., Jain, S., McCoy, A.J., et al. (2019). Macromolecular structure determination using X-rays, neutrons and electrons: recent developments in Phenix. Acta Crystallogr. D Struct. Biol. 75, 861–877. https://doi.org/10.1107/S2059798319011471.

Liu, H., Zhang, Q., Wei, P., Chen, Z., Aviszus, K., Yang, J., Downing, W., Jiang, C., Liang, B., Reynoso, L., et al. (2021). The basis of a more contagious 501Y.V1 variant of SARS-CoV-2. Cell Res. 31, 720–722. https://doi.org/10.1038/s41422-021-00496-8.

Liu, L., Wang, P., Nair, M.S., Yu, J., Rapp, M., Wang, Q., Luo, Y., Chan, J.F.-W., Sahi, V., Figueroa, A., et al. (2020). Potent neutralizing antibodies against multiple epitopes on SARS-CoV-2 spike. Nature 584, 450–456. https://doi.org/10.1038/s41586-020-2571-7.

Liu, L., Iketani, S., Guo, Y., Chan, J.F.-W., Wang, M., Liu, L., Luo, Y., Chu, H., Huang, Y., Nair, M.S., et al. (2022). Striking antibody evasion manifested by the Omicron variant of SARS-CoV-2. Nature 602, 676–681. https://doi.org/10.1038/s41586-021-04388-0.

Mannar, D., Saville, J.W., Zhu, X., Srivastava, S.S., Berezuk, A.M., Zhou, S., Tuttle, K.S., Kim, A., Li, W., Dimitrov, D.S., et al. (2021). Structural analysis of receptor binding domain mutations in SARS-CoV-2 variants of concern that modulate ACE2 and antibody binding. Cell Rep. 37, 110156. https://doi.org/10.1016/j.celrep.2021.110156.

Mannar, D., Saville, J.W., Zhu, X., Srivastava, S.S., Berezuk, A.M., Tuttle, K.S., Marquez, A.C., Sekirov, I., and Subramaniam, S. (2022). SARS-CoV-2 Omicron variant: Antibody evasion and cryo-EM structure of spike protein-ACE2 complex. Science 375, 760–764. https://doi.org/10.1126/science.abn7760.

Mastronarde, D.N. (2003). Serialem: A program for automated tilt series acquisition on tecnai microscopes using prediction of specimen position. Microsc. Microanal. 9, 1182–1183. https://doi.org/10.1017/S1431927603445911.

Mastronarde, D.N. (2005). Automated electron microscope tomography using robust prediction of specimen movements. J. Struct. Biol. 152, 36–51. https://doi.org/10.1016/j.jsb.2005.07.007.

McCallum, M., Walls, A.C., Sprouse, K.R., Bowen, J.E., Rosen, L.E., Dang, H.V., De Marco, A., Franko, N., Tilles, S.W., Logue, J., et al. (2021). Molecular basis of immune evasion by the Delta and Kappa SARS-CoV-2 variants. Science 374, 1621–1626. https://doi.org/10.1126/science.abl8506.

McCallum, M., Czudnochowski, N., Rosen, L.E., Zepeda, S.K., Bowen, J.E., Walls, A.C., Hauser, K., Joshi, A., Stewart, C., Dillen, J.R., et al. (2022). Structural basis of SARS-CoV-2 Omicron immune evasion and receptor engagement. Science 375, 864–868. https://doi.org/10.1126/science.abn8652.

Mlcochova, P., Kemp, S.A., Dhar, M.S., Papa, G., Meng, B., Ferreira, I.A.T.M., Datir, R., Collier, D.A., Albecka, A., Singh, S., et al. (2021). SARS-CoV-2 B.1.617.2 Delta variant replication and immune evasion. Nature 599, 114–119. https://doi.org/10.1038/s41586-021-03944-y.

Mou, Y., Zhou, X.X., Leung, K., Martinko, A.J., Yu, J.-Y., Chen, W., and Wells, J.A. (2018). Engineering improved antiphosphotyrosine antibodies based on an immunoconvergent binding motif. J. Am. Chem. Soc. 140, 16615–16624. https://doi.org/10.1021/jacs.8b08402.

Park, Y.-J., De Marco, A., Starr, T.N., Liu, Z., Pinto, D., Walls, A.C., Zatta, F., Zepeda, S.K., Bowen, J.E., Sprouse, K.R., et al. (2022). Antibody-mediated broad sarbecovirus neutralization through ACE2 molecular mimicry. Science 375, 449–454. https://doi.org/10.1126/science.abm8143.

Pettersen, E.F., Goddard, T.D., Huang, C.C., Couch, G.S., Greenblatt, D.M., Meng, E.C., and Ferrin, T.E. (2004). UCSF Chimera—a visualization system for exploratory research and analysis. J. Comput. Chem. 25, 1605–1612. https://doi.org/10.1002/jcc.20084.

Pintilie, G., Zhang, K., Su, Z., Li, S., Schmid, M.F., and Chiu, W. (2020). Measurement of atom resolvability in cryo-EM maps with Q-scores. Nat. Methods 17, 328–334. https://doi.org/10.1038/s41592-020-0731-1.

Planas, D., Saunders, N., Maes, P., Guivel-Benhassine, F., Planchais, C., Buchrieser, J., Bolland, W.-H., Porrot, F., Staropoli, I., Lemoine, F., et al. (2022). Considerable escape of SARS-CoV-2 Omicron to antibody neutralization. Nature 602, 671–675. https://doi.org/10.1038/s41586-021-04389-z.

Punjani, A., and Fleet, D.J. (2021). 3D variability analysis: Resolving continuous flexibility and discrete heterogeneity from single particle cryo-EM. J. Struct. Biol. 213, 107702. https://doi.org/10.1016/j.jsb.2021.107702.

Punjani, A., Rubinstein, J.L., Fleet, D.J., and Brubaker, M.A. (2017). cryoSPARC: algorithms for rapid unsupervised cryo-EM structure determination. Nat. Methods 14, 290–296. https://doi.org/10.1038/nmeth.4169.

Punjani, A., Zhang, H., and Fleet, D.J. (2020). Non-uniform refinement: adaptive regularization improves single-particle cryo-EM reconstruction. Nat. Methods 17, 1214–1221. https://doi.org/10.1038/s41592-020-00990-8.

Shi, R., Shan, C., Duan, X., Chen, Z., Liu, P., Song, J., Song, T., Bi, X., Han, C., Wu, L., et al. (2020). A human neutralizing antibody targets the receptor-binding site of SARS-CoV-2. Nature 584, 120–124. https://doi.org/10.1038/s41586-020-2381-y.

Tan, Y.Z., Baldwin, P.R., Davis, J.H., Williamson, J.R., Potter, C.S., Carragher, B., and Lyumkis, D. (2017). Addressing preferred specimen orientation in single-particle cryo-EM through tilting. Nat. Methods 14, 793–796. https://doi.org/10.1038/nmeth.4347.

Tian, F., Tong, B., Sun, L., Shi, S., Zheng, B., Wang, Z., Dong, X., and Zheng, P. (2021). N501Y mutation of spike protein in SARS-CoV-2 strengthens its binding to receptor ACE2. ELife 10. https://doi.org/10.7554/eLife.69091.

Tortorici, M.A., Beltramello, M., Lempp, F.A., Pinto, D., Dang, H.V., Rosen, L.E., McCallum, M., Bowen, J., Minola, A., Jaconi, S., et al. (2020). Ultrapotent human antibodies protect against SARS-CoV-2 challenge via multiple mechanisms. Science 370, 950–957. https://doi.org/10.1126/science.abe3354.

VanBlargan, L.A., Errico, J.M., Halfmann, P., Zost, S.J., Crowe, J.E., Purcell, L.A., Kawaoka, Y., Corti, D., Fremont, D.H., and Diamond, M. (2021). An infectious SARS-CoV-2 B.1.1.529 Omicron virus escapes neutralization by several therapeutic monoclonal antibodies. BioRxiv https://doi.org/10.1101/2021.12.15.472828.

Viana, R., Moyo, S., Amoako, D.G., Tegally, H., Scheepers, C., Althaus, C.L., Anyaneji, U.J., Bester, P.A., Boni, M.F., Chand, M., et al. (2022). Rapid epidemic expansion of the SARS-CoV-2 Omicron variant in southern Africa. Nature 603, 679–686. https://doi.org/10.1038/s41586-022-04411-y.

Walls, A.C., Park, Y.-J., Tortorici, M.A., Wall, A., McGuire, A.T., and Veesler, D. (2020). Structure, Function, and Antigenicity of the SARS-CoV-2 Spike Glycoprotein. Cell 181, 281–292.e6. https://doi.org/10.1016/j.cell.2020.02.058.

Wang, R.Y.-R., Song, Y., Barad, B.A., Cheng, Y., Fraser, J.S., and DiMaio, F. (2016). Automated structure refinement of macromolecular assemblies from cryo-EM maps using Rosetta. ELife 5. https://doi.org/10.7554/eLife.17219.

Weidenbacher, P.A.-B., Waltari, E., de los Rios Kobara, I., Bell, B.N., Pak, J.E., and Kim, P.S. (2022). Converting non-neutralizing SARS-CoV-2 antibodies targeting conserved epitopes into broad-spectrum inhibitors through receptor blockade. BioRxiv https://doi.org/10.1101/2022.01.24.477625.

Weinberg, Z.Y., Hilburger, C.E., Kim, M., Cao, L., Khalid, M., Elmes, S., Diwanji, D., Hernandez, E., Lopez, J., Schaefer, K., et al. (2021). Sentinel cells enable genetic detection of SARS-CoV-2 Spike protein. BioRxiv https://doi.org/10.1101/2021.04.20.440678.

Wilhelm, A., Widera, M., Grikscheit, K., Toptan, T., Schenk, B., Pallas, C., Metzler, M., Kohmer, N., Hoehl, S., Helfritz, F.A., et al. (2021). Reduced Neutralization of SARS-CoV-2 Omicron Variant by Vaccine Sera and monoclonal antibodies. MedRxiv https://doi.org/10.1101/2021.12.07.21267432.

Wirnsberger, G., Monteil, V., Eaton, B., Postnikova, E., Murphy, M., Braunsfeld, B., Crozier, I., Kricek, F., Niederhöfer, J., Schwarzböck, A., et al. (2021). Clinical grade ACE2 as a universal agent to block SARS-CoV-2 variants. BioRxiv https://doi.org/10.1101/2021.09.10.459744.

Yan, R., Zhang, Y., Li, Y., Ye, F., Guo, Y., Xia, L., Zhong, X., Chi, X., and Zhou, Q. (2021). Structural basis for the different states of the spike protein of SARS-CoV-2 in complex with ACE2. Cell Res. 31, 717–719. https://doi.org/10.1038/s41422-021-00490-0.

Zhang, L., Narayanan, K.K., Cooper, L., Chan, K.K., Devlin, C.A., Aguhob, A., Shirley, K., Rong, L., Rehman, J., Malik, A.B., et al. (2022). An engineered ACE2 decoy receptor can be administered by inhalation and potently targets the BA.1 and BA.2 omicron variants of SARS-CoV-2. BioRxiv https://doi.org/10.1101/2022.03.28.486075.

Zheng, S.Q., Palovcak, E., Armache, J.-P., Verba, K.A., Cheng, Y., and Agard, D.A. (2017). MotionCor2: anisotropic correction of beam-induced motion for improved cryo-electron microscopy. Nat. Methods 14, 331–332. https://doi.org/10.1038/nmeth.4193.

Zhou, T., Tsybovsky, Y., Gorman, J., Rapp, M., Cerutti, G., Chuang, G.-Y., Katsamba, P.S., Sampson, J.M., Schön, A., Bimela, J., et al. (2020a). Cryo-EM Structures of SARS-CoV-2 Spike without and with ACE2 Reveal a pH-Dependent Switch to Mediate Endosomal Positioning of Receptor-Binding Domains. Cell Host Microbe 28, 867–879.e5. https://doi.org/10.1016/j.chom.2020.11.004.

Zhou, X.X., Bracken, C.J., Zhang, K., Zhou, J., Mou, Y., Wang, L., Cheng, Y., Leung, K.K., and Wells, J.A. (2020b). Targeting Phosphotyrosine in Native Proteins with Conditional, Bispecific Antibody Traps. J. Am. Chem. Soc. 142, 17703–17713. https://doi.org/10.1021/jacs.0c08458.

Zhu, X., Mannar, D., Srivastava, S.S., Berezuk, A.M., Demers, J.-P., Saville, J.W., Leopold, K., Li, W., Dimitrov, D.S., Tuttle, K.S., et al. (2021). Cryo-electron microscopy structures of the N501Y SARS-CoV-2 spike protein in complex with ACE2 and 2 potent neutralizing antibodies. PLoS Biol. 19, e3001237. https://doi.org/10.1371/journal.pbio.3001237.

Zost, S.J., Gilchuk, P., Case, J.B., Binshtein, E., Chen, R.E., Nkolola, J.P., Schäfer, A., Reidy, J.X., Trivette, A., Nargi, R.S., et al. (2020). Potently neutralizing and protective human antibodies against SARS-CoV-2. Nature 584, 443–449. https://doi.org/10.1038/s41586-020-2548-6.

