## Supplemental Information for "Computational pipeline provides mechanistic understanding of Omicron variant of concern neutralizing engineered ACE2 receptor traps"

**Figure S1.**

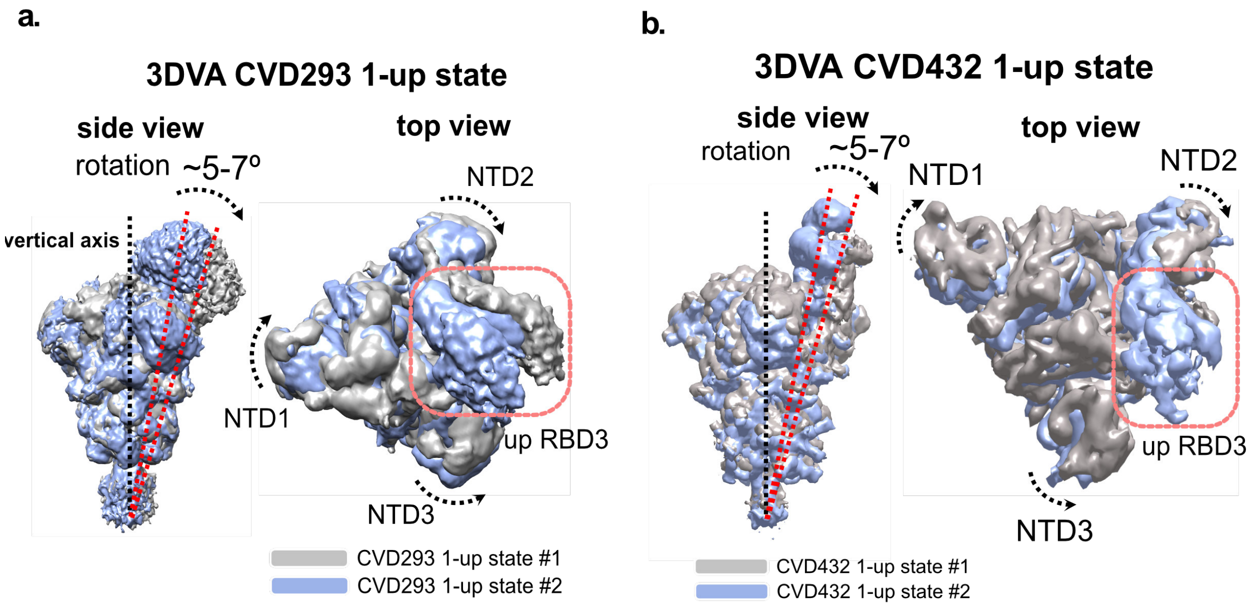

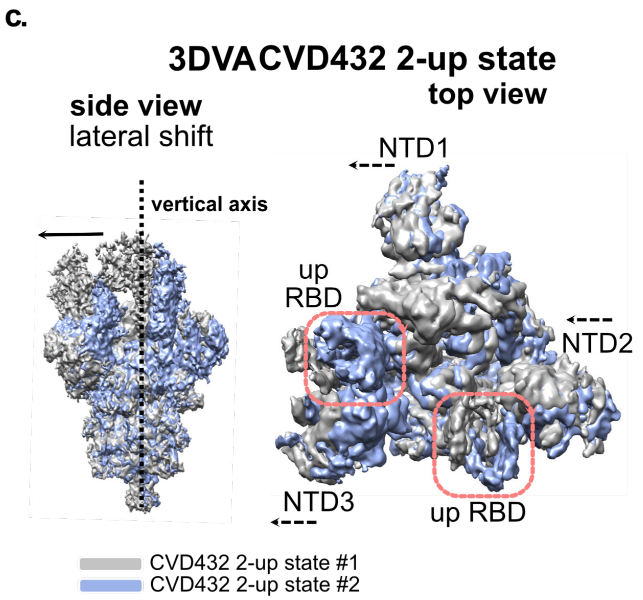

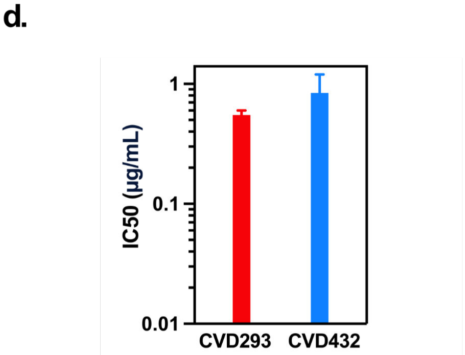

**Pseudotyped virus neutralization assay**

| ID | IC50 (µg/mL) |
|---|---|
| Spike trimer WT/CVD293 | 0.55 ± 0.05 |
| Spike trimer WT/CVD432 | 0.84 ± 0.35 |

**Figure S1. 3D variability analysis (3DVA) of WT-fl-Spike in complex with ACE2 receptor traps and Pseudovirus neutralization assay**

**a-b.** 3DVA of WT-fl-Spike/CVD293 or WT-fl-Spike/CVD432 complex in 1-RBD-up state shows ~5-7° rotation relative to the vertical axis of rotation in the RBD-ACE2 interface (side view). Also seen is the clockwise rotation of Spike-NTD1 and Spike-NTD2 while Spike-NT3 next to the 1-up-RBD undergoes a counterclockwise rotation (top view). **c.** 3DVA of WT-fl-Spike/CVD432 complex in 2-RBD-up state shows a lateral shift relative to the vertical axis of rotation of the RBDs and NTDs (top view, side view) **d.** CVD293 and CVD432 potently neutralize vesicular stomatitis virus (VSV) pseudotyped with SARS-CoV-2 WT-Spike (Wuhan-hu-1 B.1 strain with D614G mutation only).

Error bars represent standard deviation over all technical replicates from two biological replicates.

**Figure S2.**

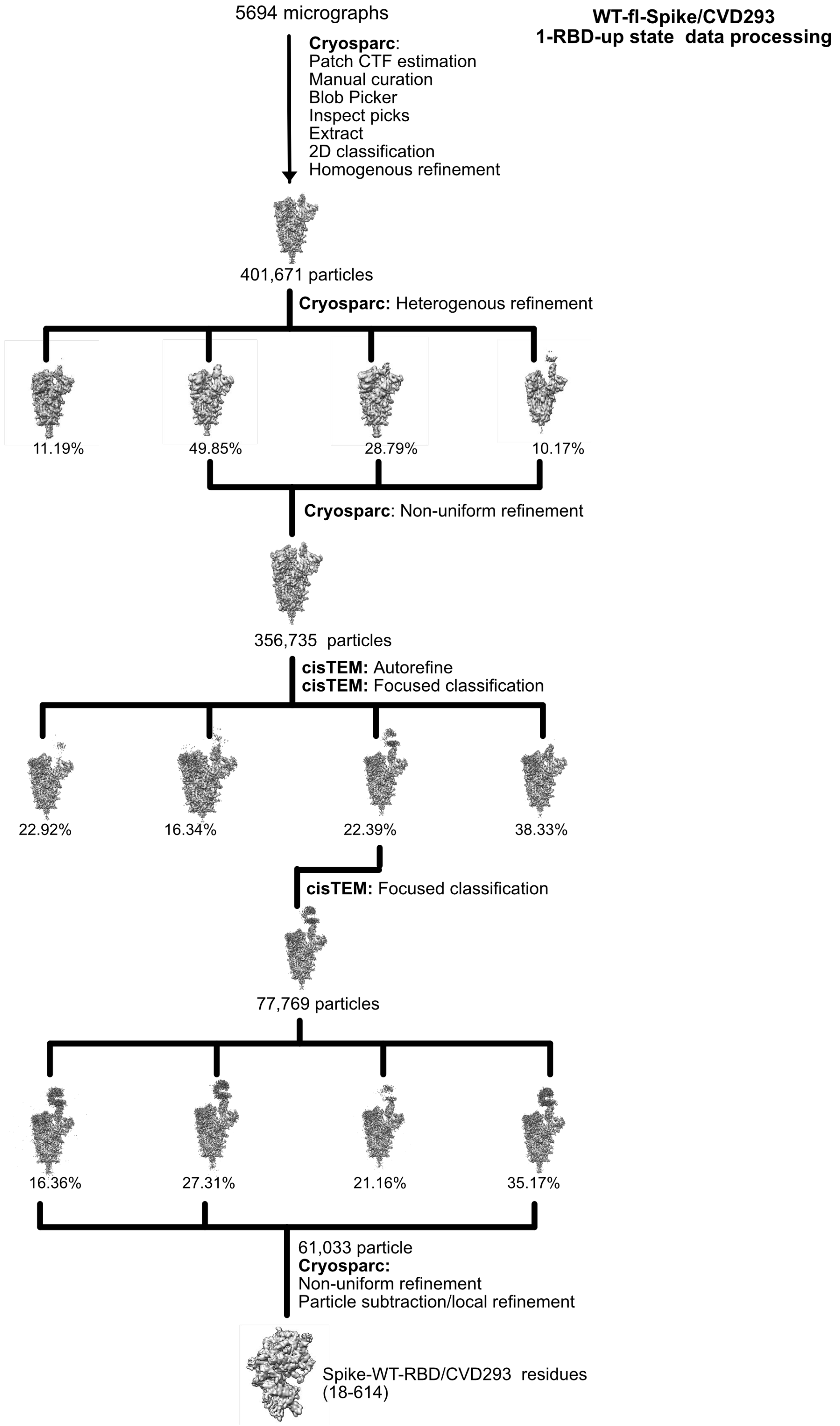

**Figure S2. Cryo-EM data processing data processing workflow for WT-fl-Spike/CVD293 1-RBD-up state**

**Figure S3.**

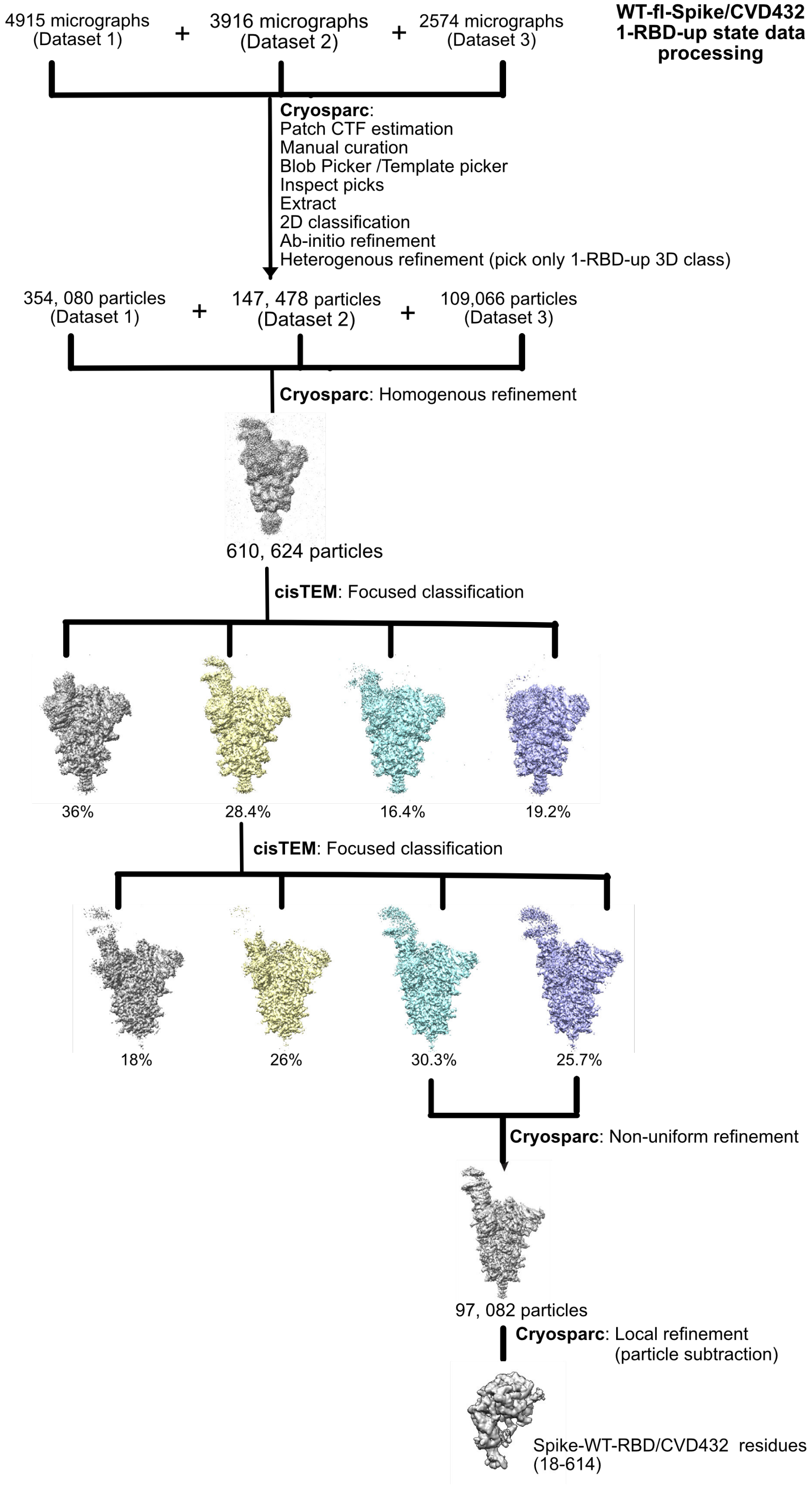

**Figure S3. Cryo-EM data processing data processing workflow for WT-fl-Spike/CVD432 1-RBD-up state**

**Figure S4.**

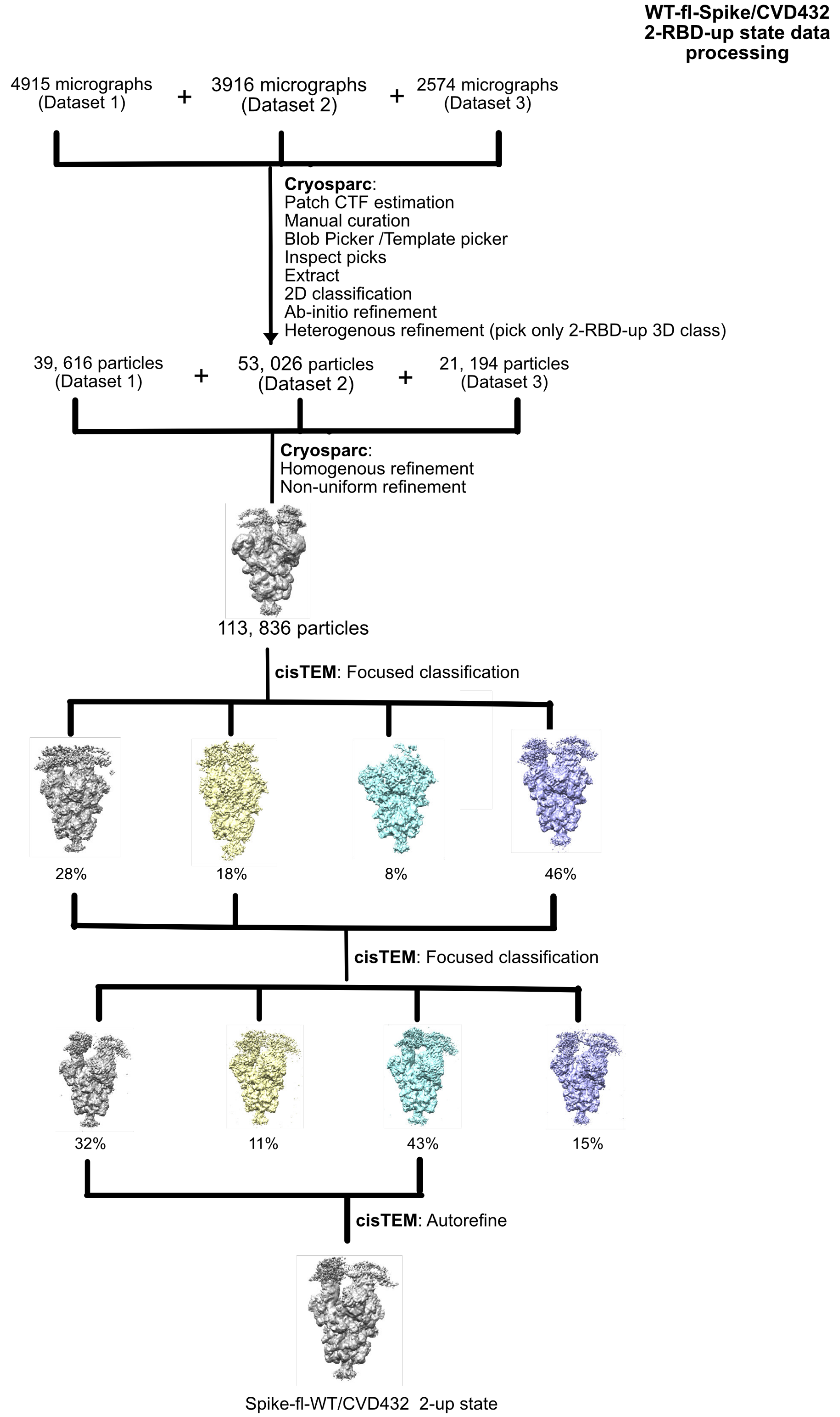

**Figure S4. Cryo-EM data processing data processing workflow for WT-fl-Spike/CVD432 2-RBD-up state**

**Figure S5.**

**a.**

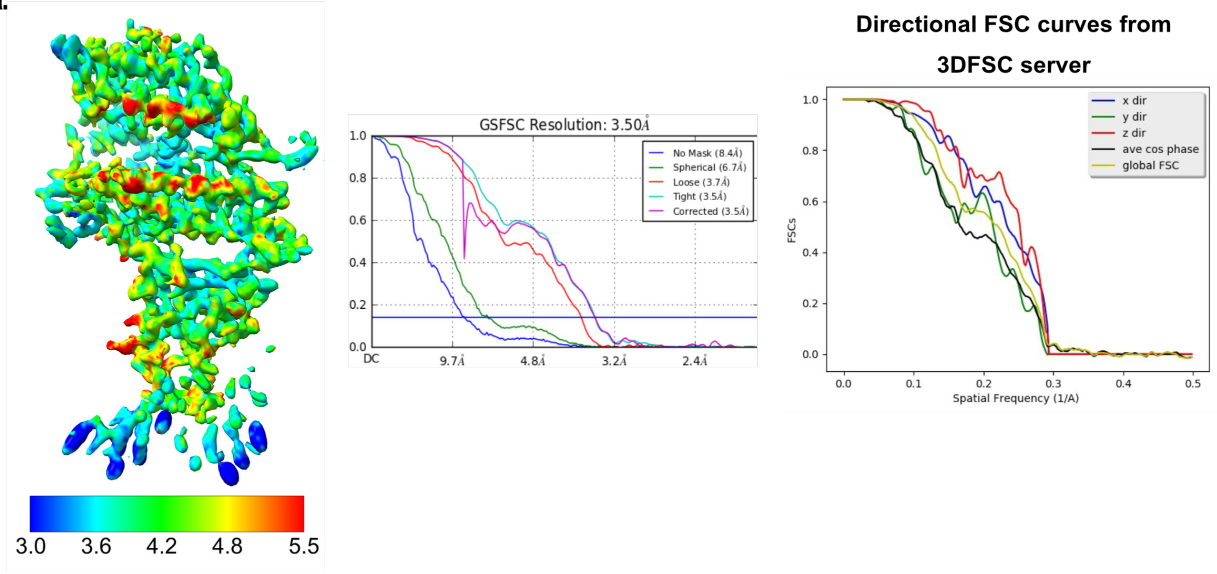

**b.**

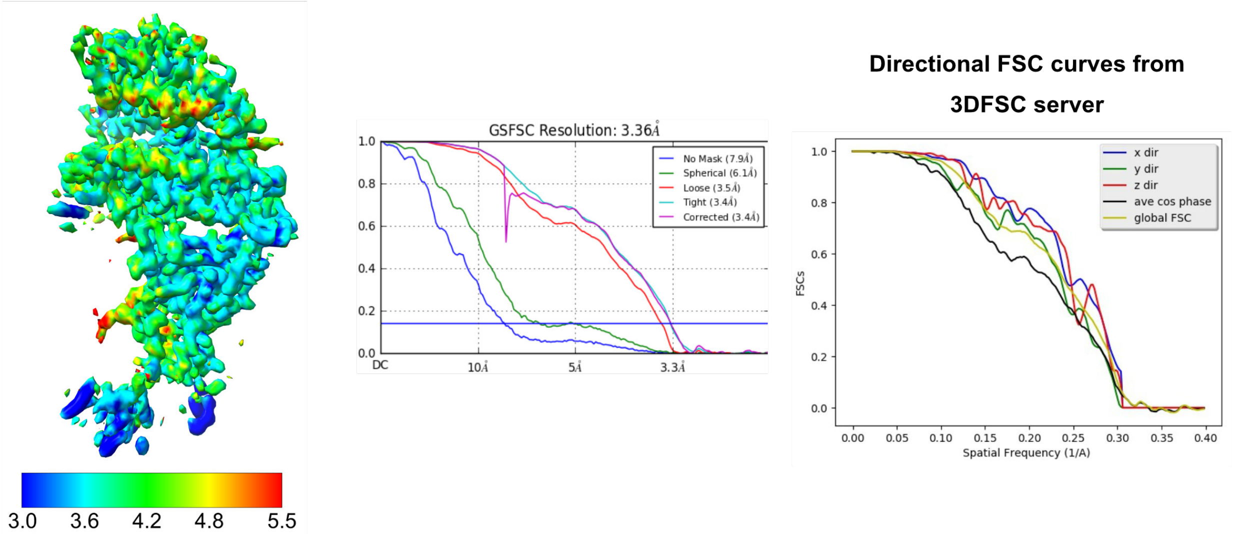

**Figure S5. Resolution estimates of the CVD293 and CVD432 cryo-EM maps.**

**a**. Left panel - Final cryo-EM reconstructions from focused refinement of WT-Spike-RBD with CVD293 colored by estimated local resolution estimated with ResMap. Middle panel - Global FSC curve for the WT-Spike-RBD/CVD293 cryo-EM reconstruction as output from cryoSPARC. In purple is the final corrected masked FSC. In blue is an un-masked FSC. In green is FSC with a soft spherical mask applied to the reconstruction. In red is FSC with loose soft solvent mask applied to the reconstruction (dilation distance of 25 Å for value of 1.0 and 40 Å for value of 0.0). In cyan is FSC with right soft solvent mask applied (dilation distance of 6 Å for value of 1.0 and 12 Å for value of 0.0). Right panel - Directional FSC curves for WT-Spike-RBD/CVD293 cryo-EM reconstruction from 3DFSC server. **b.** Left panel - Final cryo-EM reconstructions from focused refinement of WT-Spike-RBD with CVD432 colored by estimated local resolution estimated with ResMap. Middle panel – Set of global FSC curves for the WT-Spike-RBD/CVD432 cryo-EM reconstruction, colors and preparation same as in (a). Right panel - Directional FSC curves for WT-Spike-RBD/CVD432 cryo-EM reconstruction from 3DFSC server.

**Figure S6.**

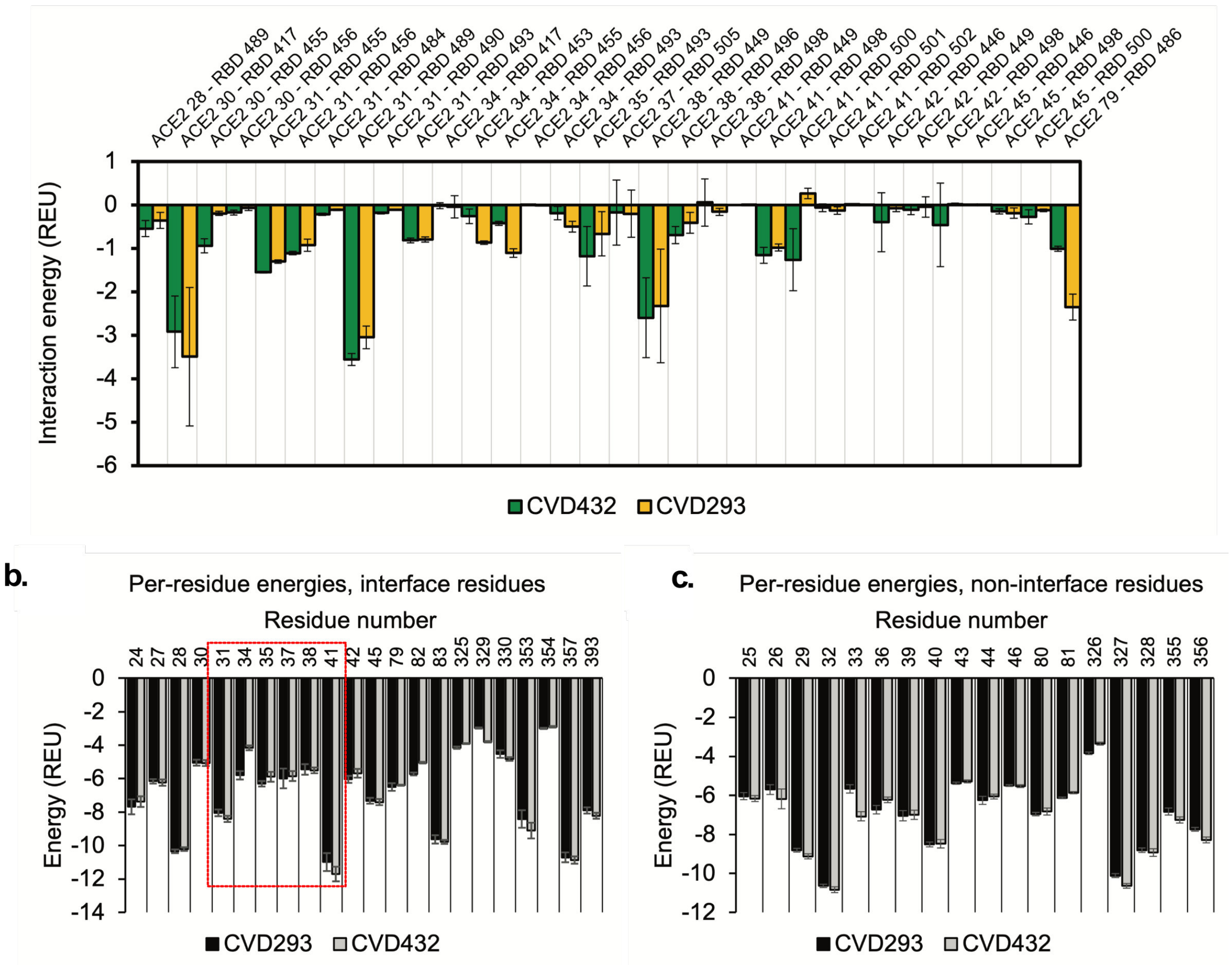

**Figure S6. Interface and total Rosetta energy calculations for model ensembles.**

**a.** Residue-level pairwise interface energies for all cryo-EM based models of CVD293/WT-Spike-RBD and CVD432/WT-Spike-RBD complexes **b-c.** Per-residue energy of CVD293/CVD432 interface helix residues with WT-Spike-RBD interface and non-interface residues.

**Figure S7.**

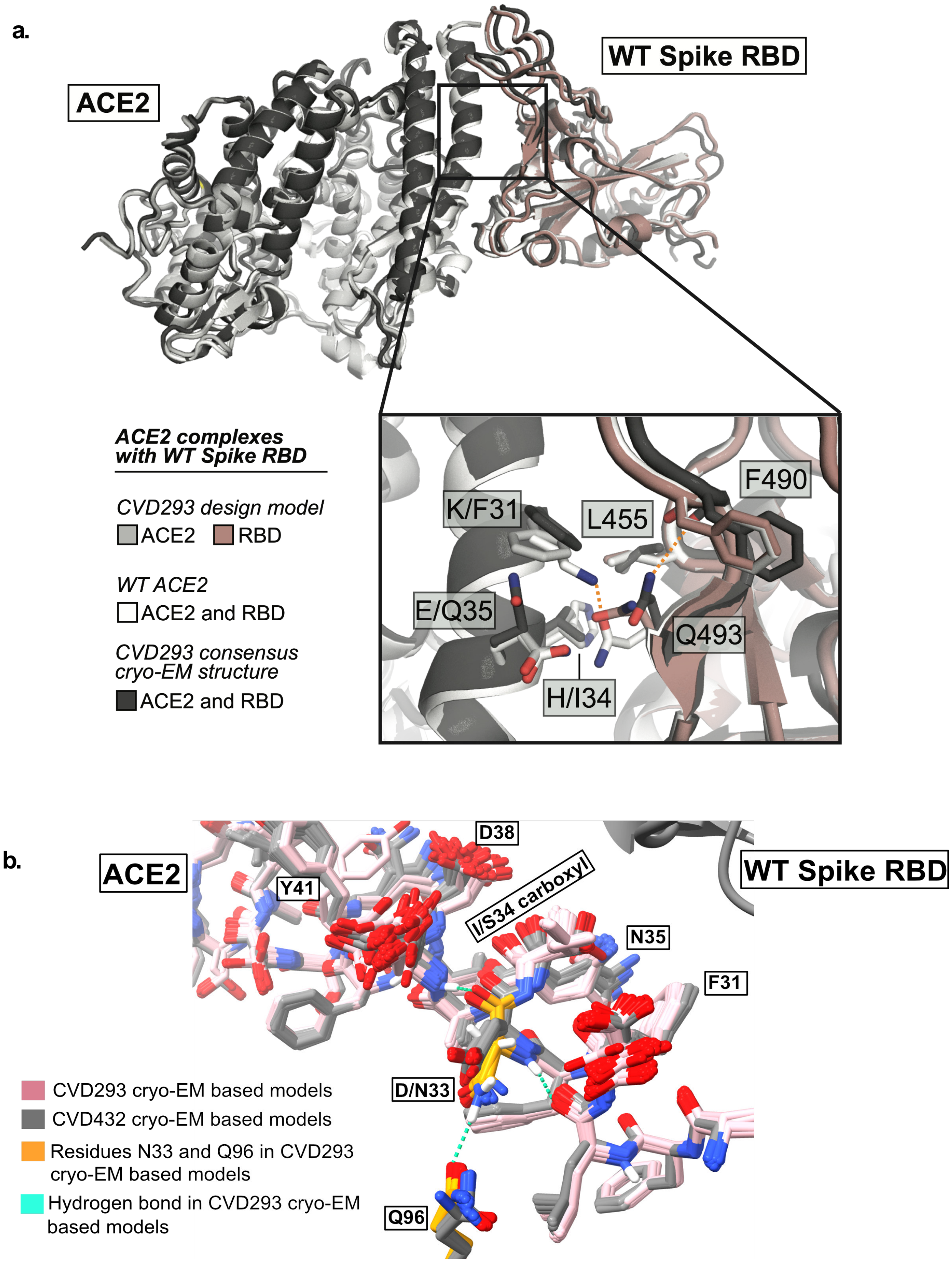

**Figure S7. Global overlays of CVD293 model between designed and experimental structure.**

**a**. Overlay of CVD293 design model and CVD293 cryo-EM “consensus” model. Inset shows the relevant interface residues comparing their rotamer positions in the complexes between CVD293 Rosetta design model and WT-ACE2 (with WT-ACE2 in gray, WT-Spike-RBD in salmon); WT-ACE2/WT-Spike-RBD (in white) and CVD293 cryo-EM “final consensus” model and WT-ACE2 (in black) **b.** Overlay of CVD293 and CVD432 cryo-EM based models showing the N33D mutation and the associated subtle shift in neighboring residues towards RBD.

**Figure S8.**

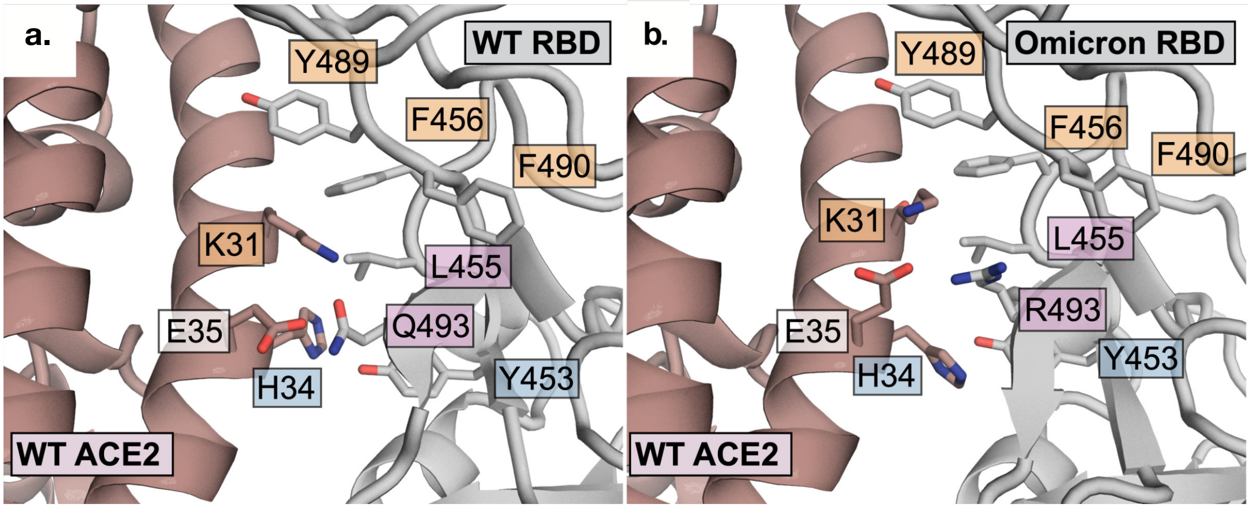

**Figure S8. Comparison of the published WT-Spike-RBD/WT-ACE2 and Omicron-RBD/WT-ACE2 a-b**. Zoomed in view of the interface of the complex WT-Spike-RBD/WT-ACE2 (PDB ID: 6M0J) and Omicron-RBD/WT-ACE2 (PDB ID: 7T9L). The structures were used to build our Omicron/CVD293 or Omicron/CVD432 models as well as for Rosetta interface based prediction of binding affinity of Omicron-RBD to our ACE2 receptor traps.

**Figure S9.**

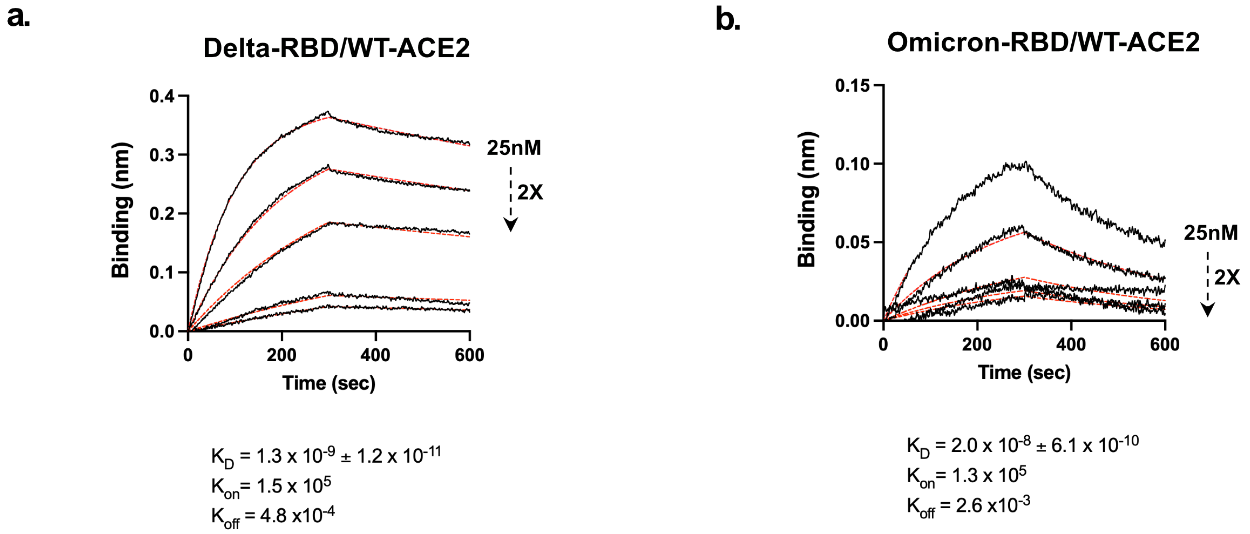

**Figure S9. Binding measurements between WT-ACE2 and Delta/Omicron-RBD.**
**a-b.** Biolayer interferometry measurements for WT-ACE2-Fc-fusion with Omicron- or Delta-RBD.

**Supplemental Table S1**

| WT-RBD/ WT-ACE2 | Omi-RBD/ WT-ACE2 | WT-RBD/ CVD293 | Omi-RBD/ CVD293 | WT-RBD/ CVD432 | Omi-RBD/ CVD432 |
|---|---|---|---|---|---|
| Q493/ E35 | R493/ E35 | Q493/ Q35 | R493/ Q35 | Q493/ Q35 | R493/ Q35 |
| Q498/Q42 | R498/Q42 | Q498/Q42 | R498/Q42 | Q498/Q42 | R498/Q42 |
| | R498/D38 | | R498/Q42 | | R498/Q42 |
| G496/K353 | S496/K353 | G496/K353 | S496/K353 | G496/K353 | S496/K353 |
| N501/Y41 | Y501/Y41 | N501/Y41 | Y501/Y41 | N501/Y41 | Y501/Y41 |
| Y505/E37 | H505/E37 | Y505/E37 | H505/E37 | Y505/E37 | H505/E37 |
| K417/D30 | N417/D30 | K417/D30 | N417/D30 | K417/D30 | N417/D30 |
| L455/K31 | L455/K31 | L455/F31 | L455/F31 | L455/F31 | L455/F31 |

**Interactions between Omicron-RBD (Omi-RBD)/WT-ACE2, CVD293 and CVD432.**
New interactions (Eg: hydrogen bonds) not present in WT-Spike-RBD/WT-ACE2 or present in Omi-RBD/WT-ACE2, Omi-RBD/CVD293 or Omi-RBD/CVD432 even with mutations in Omi-RBD are shown in green. Interactions present in WT-Spike-RBD/WT-ACE2 but not present in Omi-RBD/WT-ACE2, Omi-RBD/CVD293 or Omi-RBD/CVD432 are shown in red

**Supplementary Table S2. Apparent binding affinities and associated $k_{on}$ and $k_{off}$ of ACE2 receptors traps or WT-ACE2- Fc-fusion for WT-Spike-RBD, Delta-RBD and Omicron-RBD.**

| | $K_D$ (M) | $k_{on}$ (1/Ms) | $k_{off}$ (1/s) |
|---|---|---|---|
| **WT-Spike-RBD/WT-ACE2-Fc-fusion** | $3.1x10^{-9}$ ±$3.4x10^{-11}$ | $3.1 \times 10^{5}$ | $9.6 \times 10^{-4}$ |
| **Delta-RBD/WT-ACE2-Fc-fusion** | $1.3x10^{-9}$ ±$1.2x10^{-11}$ | $1.5 \times 10^{5}$ | $4.8 \times 10^{-4}$ |
| **Omicron-RBD/WT-ACE2-Fc-fusion** | $2.0x10^{-8}$ ±$6.1x10^{-10}$ | $1.3 \times 10^{5}$ | $2.6 \times 10^{-3}$ |
| **WT-Spike-RBD/CVD293** | $5.1x10^{-9}$ ±$6.1x10^{-11}$ | $4.2 \times 10^{5}$ | $2.1 \times 10^{-3}$ |
| **Delta-RBD/CVD293** | $1.9x10^{-9}$ ±$2.2x10^{-11}$ | $9.7 \times 10^{5}$ | $1.2 \times 10^{-5}$ |
| **Omicron-RBD/CVD293** | $4.2x10^{-9}$ ±$7.4x10^{-11}$ | $2.3 \times 10^{5}$ | $1.1 \times 10^{-5}$ |
| **WT-Spike-RBD/CVD432** | $2.3x10^{-10}$ ±$7.7x10^{-12}$ | $9.3 \times 10^{5}$ | $2.1 \times 10^{-4}$ |
| **Delta-RBD/CVD432** | $7.1x10^{-11}$ ±$7.7x10^{-12}$ | $1.4 \times 10^{6}$ | $9.7 \times 10^{-5}$ |
| **Omicron-RBD/CVD432** | $4.0x10^{-9}$ ±$7.6x10^{-11}$ | $6.6 \times 10^{5}$ | $2.6x10^{-3}$ |

**Supplementary Table S3. CryoEM collection, refinement and resulting model statistics.**

| | **Spike-RBD/ CVD293** | **Spike-RBD/ CVD432** | **WT-fl-Spike/ CVD293** (1-RBD-up state) | **WT-fl-Spike/ CVD432** (1-RBD-up state**)** | **WT-fl-Spike/ CVD432** (2-RBD-up state) |
|---|---|---|---|---|---|
| **Data collection and processing** | | | | | |
| Microscope and camera | Titan Krios, K3 | | | | |
| Magnification | 105,000 | | | | |
| Voltage (kV) | 300 | | | | |
| Total dose ($e^-$/Å$^2$) | 68 | | | | |
| Dose rate ($e^-$ /physical pixel/sec) | 8 | | | | |
| Exposure per frame (sec) | 0.05 | | | | |
| Defocus range (µm) | -0.8 to -1.8 | | | | |
| Physical pixel size (Å) | 0.834 | | | | |
| Symmetry imposed | C1 | | | | |
| Initial particle images (no.) | 2,505,078 | 2,794,745 | 2,505,078 | 2,794,745 | 3,847,252 |
| Final particle images (no.) | 61,033 | 97,082 | 61,033 | 97,082 | 97,374 |
| Map resolution (Å) | 3.5 | 3.36 | 3.77 | 3.5 | 2.97 |
| FSC threshold | 0.143 | 0.143 | 0.143 | 0.143 | 0.143 |
| Map resolution range (Å) | 3-15 | 3-15 | 2-12 | 2-20 | 2-20 |
| **Refinement** | | | | | |
| Initial model used (PDB code) | 7DX5, 6M0J | 7DX5, 6M0J | | | |
| Map/Model FSC (Å) | 4.0 | 3.7 | | | |
| FSC threshold | 0.5 | 0.5 | | | |
| Map sharpening *B* factor (Å2) | -62.6 | -53.7 | | | |

| | | |
|---|---|---|
| Map/Model cross correlation, masked | 0.77 | 0.76 |
| **Model composition** | | |
| Non-hydrogen atom**s** | 6,447 | 6,445 |
| Protein residues | 796 | 796 |
| Ligands | 0 | 0 |
| **B factors ($A^2$)** | | |
| Protein | 100.3 | 88.4 |
| **R.M.S. deviations** | | |
| Bond lengths (Å) | 0.010 | 0.011 |
| Bond angles (°) | 1.137 | 1.493 |
| **Validation** | | |
| MolProbity score | 0.96 | 0.83 |
| Clashscore | 0.87 | 0.48 |
| Poor rotamers (%) | 0.00 | 0.14 |
| **Ramachandran plot** | | |
| Favored (%) | 96.84 | 97.10 |
| Allowed (%) | 3.03 | 2.53 |
| Disallowed (%) | 0.13 | 0.38 |

### STAR ★ METHODS

### KEY RESOURCE TABLE

| REAGENT or RESOURCE | SOURCE | IDENTIFIER |
|---|---|---|
| **Pseudovirus strains** | | |
| WT-Spike (D614G) (based on SARS-CoV-2 B.1 strain) | This study | WT-Spike pseudovirus |
| Delta-Spike (based on SARS-CoV-2 B.1.617.2 strain) | This study | Delta-Spike pseudovirus |
| Omicron-Spike (based on SARS-CoV-2 B.1.1.529 strain) | This study | Omicron-Spike pseudovirus |
| **Chemicals, peptides, and recombinant proteins** | | |
| WT-fl-Spike | This study | WT-fl-Spike |
| ACE2 computationally engineered protein | This study | CVD293 |
| ACE2 affinity matured protein | This study | CVD313 |
| ACE2 linker variant of CVD313 | This study | CVD432 |
| WT-Spike-RBD | This study | WT-Spike-RBD |
| Delta-Spike-RBD | This study | Delta-Spike-RBD |
| Omicron-Spike-RBD | This study | Omicron-Spike-RBD |
| **Deposited data** | | |
| Spike-RBD/CVD293 | EMDB - (https://www.ebi.ac.uk/emdb/); PDB - (https://www.rcsb.org) | |

| | | |
|---|---|---|
| Spike-RBD/CVD432 | EMDB - (https://www.ebi.ac.uk/emdb/); PDB - (https://www.rcsb.org) | |
| WT-fl-Spike/CVD293 (1-RBD-up state) | EMDB - (https://www.ebi.ac.uk/emdb/); PDB - (https://www.rcsb.org) | |
| WT-fl-Spike/CVD432 (1-RBD-up state) | EMDB - (https://www.ebi.ac.uk/emdb/); PDB - (https://www.rcsb.org) | |
| WT-fl-Spike/CVD432 (2-RBD-up state) | EMDB - (https://www.ebi.ac.uk/emdb/); PDB - (https://www.rcsb.org) | |
| **Oligonucleotides** | | |
| WT-fl-Spike | Gift from Pak lab and Krammer lab | WT-fl-Spike |
| ACE2 computationally engineered protein | (Glasgow et al., 2020) | CVD293 |
| ACE2 affinity matured protein | (Glasgow et al., 2020) | CVD313 |
| ACE2 linker variant of CVD313 | Twist Biosciences | CVD432 |
| WT-Spike-RBD | (Glasgow et al., 2020) | WT-Spike-RBD |
| Delta-Spike-RBD | This study | Delta-Spike-RBD |
| Omicron-Spike-RBD | This study | Omicron-Spike-RBD |
| **Software and algorithms** | | |
| cryoSPARC | (Punjani et al., 2017) | https://cryosparc.com |
| MotionCor2 | (Zheng et al., 2017) | https://docs.google.com/forms/d/e/1FAIpQLSfAQm5MA81qTx90W9JL6ClzSrM77tytsvyyHh1ZZWrFByhmfQ/viewform |

| | | |
|---|---|---|
| SerialEM | (Mastronarde, 2003, 2005) | https://bio3d.colorado.edu/SerialEM/ |
| cisTEM | (Grant et al., 2018) | https://bhimes.github.io/cisTEM_docs/intro.html |
| CHIMERA | (Pettersen et al., 2004) | https://www.cgl.ucsf.edu/chimera/ |
| Phenix Real Space Refine | (Liebschner et al., 2019) | https://phenix-online.org |
| Rosetta (2020.08 release) | (Wang et al., 2016) | For automated structure refinement |
| Rosetta version 2021.48.post.dev+8.master.77491fa20be 77491fa20be83588cfc37ab422ba5b95eca128eb git@github.com:RosettaCommons/main.git 2021-12-02T08:56:13 | (Alford et al., 2017) (Leman et al., 2020) | Protein design and structural analysis |
| COOT 0.9 | (Emsley et al., 2010) | https://www2.mrc-lmb.cam.ac.uk/personal/pemsley/coot/ |
| ISOLDE 1.0 | (Croll, 2018) | |
| ResMap | (Kucukelbir et al., 2014) | http://resmap.sourceforge.net |
| 3DFSC server | (Tan et al., 2017) | https://3dfsc.salk.edu |
| Q-scores | (Pintilie et al., 2020) | https://github.com/gregdp/mapq |
| FlowJo™ | | https://www.flowjo.com |
| Octet Data Analysis HT software version 10.0 | | https://www.sartorius.com/en/products/protein-analysis/octet-systems-software |
| GraphPad Prism 9 Version 9.3.0 (345) | | https://www.graphpad.com/scientific-software/prism/ |
